# A *Plasmodium falciparum* MORC protein complex modulates epigenetic control of gene expression through interaction with heterochromatin

**DOI:** 10.1101/2023.09.11.557196

**Authors:** Maneesh Kumar Singh, Victoria A. Bonnell, Israel Tojal Da Silva, Verônica Feijoli Santiago, Miriam S. Moraes, Jack Adderley, Christian Doerig, Giuseppe Palmisano, Manuel Llinás, Célia R. S. Garcia

## Abstract

Dynamic control of gene expression is critical for blood stage development of malaria parasites. Here, we used multi-omic analyses to investigate transcriptional regulation by the chromatin-associated microrchidia protein, MORC, during asexual blood stage development of the human malaria parasite *Plasmodium falciparum*. We show that *Pf*MORC (PF3D7_1468100) interacts with a suite of nuclear proteins, including APETALA2 (AP2) transcription factors (*Pf*AP2-G5, *Pf*AP2-O5, *Pf*AP2-I, PF3D7_0420300, PF3D7_0613800, PF3D7_1107800, and PF3D7_1239200), a DNA helicase DS60 (PF3D7_1227100), and other chromatin remodelers (*Pf*CHD1 and *Pf*EELM2). Transcriptomic analysis of *Pf*MORC^HA-glmS^ knockdown parasites revealed 163 differentially expressed genes belonging to hypervariable multigene families, along with upregulation of genes mostly involved in host cell invasion. *In vivo* genome-wide chromatin occupancy analysis during both trophozoite and schizont stages of development demonstrates that *Pf*MORC is recruited to repressed, multigene families, including the *var* genes in subtelomeric chromosomal regions. Collectively, we find that *Pf*MORC is found in chromatin complexes that play a role in the epigenetic control of asexual blood stage transcriptional regulation and chromatin organization.

## Introduction

Despite global efforts to combat malaria, the disease caused an estimated 249 million new cases and more than 608,000 deaths in 2022 (World Malaria Report 2023). The etiological agents of human malaria are apicomplexan parasites of the genus *Plasmodium*. Among the six known *Plasmodium* species that can infect humans, *Plasmodium falciparum* is the most lethal, causing the majority of annual deaths (Cowman *et al*. 2012). The intraerythrocytic developmental cycle (IDC) of *P. falciparum* is responsible for the clinical symptoms of malaria. The IDC commences when merozoites generated during the liver-stage enter the circulatory system and invade red blood cells (RBCs). While growing inside RBCs, parasites undergo multiple developmental phases with distinct morphological characteristics (ring, trophozoite, and schizont). Maturation and schizogony leads to the formation of 16-32 daughter merozoites destined to invade new RBCs (Singh *et al*. 2010). A small fraction (<10%) of parasites differentiate into non-replicative sexual gametocytes, which are transmitted to the mosquito during a second blood meal to complete sexual development.

Parasite development through the asexual blood stage is facilitated by precise transcriptional regulation, where genes are only expressed when needed in a just-in-time fashion (Bozdech *et al*. 2003). Approximately 90% of genes across the *P. falciparum* genome are transcribed during the asexual blood stage in a cascade of gene expression believed to be controlled by both sequence-specific transcription factors and dynamic epigenetic changes to chromatin (Painter *et al*. 2011, Toenhake *et al*. 2018, Jeninga *et al*. 2019, Hollin *et al*. 2021). The gene regulatory toolbox in malaria parasites is lacking many transcriptional regulatory factors conserved in other eukaryotes (Gardner *et al*. 2002). The identification of a large family of *Plasmodium* homologues of the plant APETALA2/Ethylene Response Factor (AP2/ERF) transcription factors (TFs) provided a breakthrough to unravel key regulators of gene expression in these parasites (Balaji *et al*. 2005). There are 28 putative APETALA2 (ApiAP2) TFs identified in *P. falciparum,* each protein containing one to three AP2 DNA binding domains (Painter *et al*. 2011, Jeninga *et al*. 2019). Multiple studies in *Plasmodium spp.,* using a variety of approaches, have characterized essential functions of ApiAP2 TFs in RBC invasion, gametocytogenesis, oocyst formation and sporozoite formation (Yuda *et al*. 2009, Kafsack *et al*. 2014, Sinha *et al*. 2014, Modrzynska *et al*. 2017, Santos *et al*. 2017, Zhang *et al*. 2018). ApiAP2 proteins in *Plasmodium* species display a wide array of functions during parasite development in both the vertebrate host and the mosquito vector, but surprisingly, over 70% of ApiAP2 proteins are expressed in the asexual blood stage of the lifecycle (Bozdech *et al*. 2003, Le Roch *et al*. 2003, Chappell *et al*. 2020).

Despite increasing evidence detailing ApiAP2 protein regulatory complexes (Santos *et al*. 2017, Harris *et al*. 2019, Hillier *et al*. 2019, Hoeijmakers *et al*. 2019, Farhat *et al*. 2020, Josling *et al*. 2020, Miao *et al*. 2021, Srivastava *et al*. 2023, Yuda *et al*. 2023, Antunes *et al*. 2024), the functional properties and specific interaction partners of many ApiAP2 TFs remain to be elucidated. A quantitative mass spectrometry-based analysis of the parasite protein interaction network has revealed links between ApiAP2 TFs and many chromatin remodelers, such as an extended Egl-27 and MTA1 homology 2 (EELM2) domain-containing proteins (PF3D7_0519800 and PF3D7_1141800), histone deacetylase protein 1 (HDAC1; PF3D7_0925700), and the microrchidia family protein *Pf*MORC (PF3D7_1468100) (Hillier *et al*. 2019). Genome-wide mutagenesis studies revealed that several genes within these proposed networks, including *pfmorc*, are essential for parasite proliferation (Bushell *et al*. 2017, Zhang *et al*. 2018). Using a targeted deletion strategy, we previously were unable to delete *pfmorc,* further suggesting that it is essential for parasite growth (Singh *et al*. 2021). Moreover, STRING network analysis has shown that the putative ApiAP2:*Pf*MORC complex forms a network with proteins having DNA-binding or nucleosome assembly properties, suggesting that *Pf*MORC may function as an accessory protein in epigenetic regulation (Hillier *et al*. 2019). Previous work identified *Pf*MORC in a chromatin complex containing the chromatin remodeling protein *Pf*ISWI (PF3D7_0624600) located at *var* gene promoter regions (Bryant *et al*. 2020). All *P. falciparum* strains encode roughly 60 highly polymorphic *var* genes, but through an epigenetic allelic exclusion mechanism, each parasite is thought to expresses a single allele (Real *et al*. 2022, Schneider *et al*. 2023). Bryant *et al*. have proposed that *Pf*MORC is recruited to these heterochromatic regions to assist in the silencing of *var* genes, on the basis of its known function as a repressor complex component in model eukaryotes. In the related apicomplexan parasite *Toxoplasma gondii*, *Tg*MORC functions as a repressor of sex-associated genes by recruiting the *Tg*HDAC3 histone deacetylase and forming heterogenous complexes with eleven ApiAP2 TFs (Farhat *et al*. 2020). To date, only two *Tg*MORC:HDAC3 complexes have been characterized, including a dimeric *Tg*AP2XII-2:HDAC3 complex and a heterotrimeric *Tg*AP2XII-1:AP2XI-2:HDAC3 complex (Srivastava *et al*. 2023, Antunes *et al*. 2024).

MORC proteins canonically consist of two major conserved regions: (1) a catalytic ATPase domain at the N-terminus (comprising a GHKL [Gyrase, HSP90, Histidine kinase, and MutL] domain and S5 fold domain) and (2) a C-terminal protein-protein interaction domain containing one or more coiled-coils (Koch *et al*. 2017). The conserved MORC gene family is present in most eukaryotes (with the exception of fungi), often with multiple paralogs per genome (Dong *et al*. 2018), and has been extensively investigated in various model systems in the context of epigenetic gene regulation. In plants, MORC proteins function in gene repression and heterochromatin compaction (Koch *et al*. 2017, Zhong *et al*. 2023). Additionally, MORC proteins have been shown to play diverse roles in metazoans by forming protein-protein interactions with immune-responsive proteins, SWI chromatin remodeling complexes, histone deacetylases, and histone tail modifications (Iyer *et al*. 2008, Kang *et al*. 2012, Moissiard *et al*. 2012, Bordiya *et al*. 2016, Kim *et al*. 2019).

While most metazoans possess five to seven MORC paralogs, apicomplexan parasites contain a single *morc* gene (Iyer *et al*. 2008), which encodes not only the canonical animal-like GHKL ATPase domain, but also three Kelch-repeats, and a CW-type zinc-finger domain not found in mammalian MORCs (Farhat *et al*. 2020). This unique domain architecture suggests that the apicomplexan MORC proteins may have parasite-specific roles. To dissect the functional roles of *Pf*MORC, we conducted a proteomic analysis with *Pf*MORC^GFP^, which identified several nuclear proteins, including a cluster of ApiAP2 TFs, as possible interacting partners. We also determined the genome-wide localization of *Pf*MORC at multiple developmental stages, which revealed *Pf*MORC recruitment predominantly to subtelomeric regions, corroborating that *Pf*MORC may act as a repressor of the clonally variant gene families that are important contributors to malaria pathogenesis. Finally, we performed transcriptomic analysis in *Pf*MORC^HA-glmS^ knockdown parasites at the asexual stage to investigate alterations in global gene expression. We observed an overrepresentation of downregulated genes belonging to the heterochromatin-associated hypervariable gene family proteins. Overall, this study allows us to assign a role for *Pf*MORC in facilitating the plasticity of epigenetic regulation during asexual blood stage development.

## Results

### Proteins that co-purify with PMORC represent gene regulatory and chromatin remodeling components

A previous study using Blue-Native PAGE identified a *Pf*MORC complex in association with ApiAP2 proteins and chromatin remodeling machinery (Hillier *et al*. 2019). To validate this observation and expand the repertoire of *Pf*MORC interactors, we used a targeted immunoprecipitation approach coupled to LC‒MS/MS proteomic quantification. We used a previously generated *Pf*MORC^GFP^ parasite line (Singh *et al*. 2021) to carry out immunoprecipitation with an anti-GFP antibody at the trophozoite stage, where *Pf*MORC is abundant (Singh *et al*. 2021). The *P. falciparum* 3D7 strain expressing wild-type *pfmorc* was used as a negative control. Trophozoite lysates were incubated with anti-GFP-Trap-A beads (ChromoTek, gta-20), and the immunocaptured proteins were resolved by SDS‒PAGE (**Supplemental Figure 1A**). We applied a label-free quantitative proteomics approach with a false discovery rate (FDR) of 1% and number of peptides ≥2 to excised gel samples to identify proteins interacting with *Pf*MORC^GFP^. From three biological replicates, we identified 211, 617, and 656 proteins respectively. We further identified the overlap between all three wild-type 3D7 and *Pf*MORC^GFP^ replicates and found a total of 132 and 142 proteins, respectively (**Supplemental Figure 1B-D**, **Supplemental Table 1**).

To analyze the relative ratio of proteins between wild-type 3D7 and *Pf*MORC^GFP^ groups, we used the mean-normalized MS/MS count to calculate a fold change from *Pf*MORC^GFP^/3D7, and selected differentially abundant proteins above a 1.5x cutoff filter. This high stringency threshold was used to preclude any mis-identification of *Pf*MORC interactors caused by variability between the replicates **(Figure 1A, Supplemental Figure 1D)**. This analysis resulted in 143 significantly enriched proteins (**Supplemental Table 2**). From these candidate PfMORC-interacting proteins, the top enriched protein (20.8-fold enrichment) was *Pf*EELM2 (PF3D7_0519800, *-log10 p-value* 3.43). *Pf*EELM2 was previously predicted as a *Pf*MORC interactor (Hillier *et al*. 2019) and identified in a quantitative histone peptide pulldown to be consistently recruited to H2B.Z_K13/14/18a (Hoeijmakers *et al*. 2019). Similarly, EELM2 of the related Apicomplexan parasite *T. gondii* was recently identified in a *Tg*MORC-associated complex (Farhat *et al*. 2020). We also detected numerous ApiAP2 transcription factors [*Pf*AP2-G5, *Pf*AP2-O5, *Pf*AP2-I, PF3D7_1107800, PF3D7_0613800, PF3D7_0420300 and PF3D7_1239200] (**Table 1**), similar to results reported both by Hillier et al. (Hillier *et al*. 2019) and in the *Toxoplasma* studies which also predicted or experimentally identified many ApiAP2 interactions (Farhat *et al*. 2020, Srivastava *et al*. 2023, Antunes *et al*. 2024). *Pf*AP2-G5 (PF3D7_1139300, *-log10 p-value* 0.22) and *Pf*AP2-O5 (PF3D7_1449500, *-log10 p-value* 0.41) were enriched 20.5-fold and 14.99-fold, respectively, suggesting that these factors are likely major components in complex with *Pf*MORC. To corroborate our results, we compared our *Pf*MORC^GFP^ coIPed proteins to a recently published, computationally predicted, protein‒protein interaction network (Hillier *et al*. 2019, Bryant *et al*. 2020, Subudhi *et al*. 2023) and found many of ApiAP2 and EELM2 proteins shared across both datasets (**Figure 1B**). Collectively, our results demonstrate a direct association of *Pf*MORC with various chromatin-associated factors, including at least seven ApiAP2 proteins. The potential interactors of *Pf*MORC that we detected in this experiment also included proteins implicated in DNA replication and repair, including the ATP-dependent RNA helicase DBP5 (PF3D7_1459000, *-log10 p-value* 0.46) and the DNA helicase 60 DH60 (PF3D7_1227100, *- log10 p-value* 0.41). *Pf*DH60 exhibits DNA and RNA unwinding activities, and its high expression in the trophozoites suggest a role in DNA replication (Pradhan *et al*. 2005). We also identified two putative chromatin-associated proteins, chromodomain-helicase-DNA-binding protein 1 CHD1 (PF3D7_1023900, *-log10 p-value* 0.30) and the SNF2 chromatin-remodeling ATPase ISWI (PF3D7_0624600, *-log10 p-value* 0.001), which are associated with chromosome structure maintenance, DNA replication, DNA repair, and transcription regulation. *Pf*ISWI was previously reported to be associated with *Pf*MORC in the context of *var* gene transcriptional activation during ring stage development (Bryant *et al*. 2020). Gene Ontology (GO) analysis was performed to identify enriched biological processes, cellular components, and molecular functions using a *p-value* cut-off of 0.05. We found significant enrichment of DNA-binding transcription factor activity and mRNA binding, transcription, and regulation of transcription activity (**Supplemental Figure 1E**, **Supplemental Table 3)**. Overall, we again find that *Pf*MORC forms a link between ApiAP2 TFs and chromatin remodelers (**Figure 1C**).

**Figure 1.**
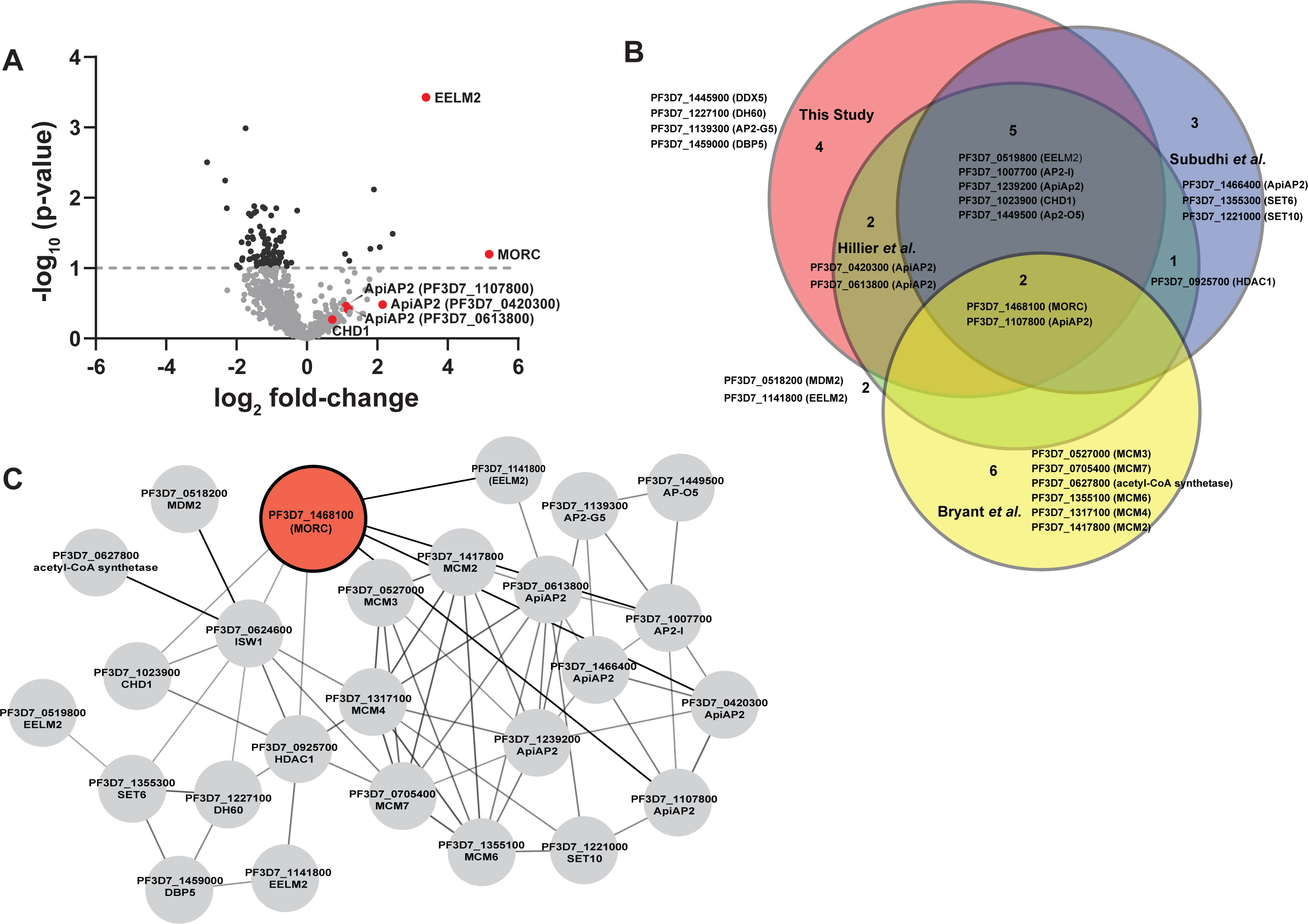
Proteomic analysis of parasites expressing *Pf*MORC^GFP^ reveals *Pf*MORC association with nuclear proteins of epigenetic regulation. **(A)** Volcano plot illustrates the protein enrichment in lable free LC-MS/MS analysis of *Pf*MORC CoIPed proteins from three independent experiments at 32hpi. For normalized MS/MS counts, student’s t-test was performed and proteins were ranked as -log2 fold-change (x-axis) versus statistical *p-values* (y-axis). Gray dashed horizontal line shows the *p-value* cutoff. **(B)** Comparative analysis showing the juxtaposition of specific proteins CoIPed in PfMORC^GFP^ with selected proteins from recent works of Hillier *et al*., Bryant *et al*. and Subudhi *et al*. where ApiAP2 or ISW1 were used as bait in similar CoIP experiments. The Venn diagram illustrates the overlap between identified proteins, revealing that the intersecting proteins are primarily ApiAP2 and chromatin remodellers. **(C)** An interactive protein-protein interaction network is constructed with proteins known to interact with PfMORC, using proteins identified in this study and proteins documented in previously published works. Proteins identified in this study with known interaction networks from the STRING database were used to curate the network employing Cytoscape to enrich the network quality.

**Table 1.** –. Potential *Pf*MORC interacting proteins enriched in CoIP eluates and identified in LC‒MS/MS from three independent experiments and fold change ≥ 1.5x GFP/3D7.

| Protein ID | Annotation | Fold Change | *-log p-value* | Known Function |
| --- | --- | --- | --- | --- |
| PF3D7_0519800 | EELM2 domain-containing protein | 20.79 | 3.45 | - |
| PF3D7_1139300 | AP2 domain transcription factor AP2-G5 | 20.5 | 0.22 | Repressor of Commitment and Early Gametocyte Development (Shang *et al.* 2021) |
| PF3D7_1449500 | AP2 domain transcription factor AP2-O5 | 14.99 | 0.41 | Regulator of Mature Ookinete Motility (Modrzynska *et al.* 2017) |
| PF3D7_1468100 | *Pf*MORC | 11.76 | 1.20 |  |
| PF3D7_1023900 | SNF2 helicase, putative or Chromodomain-helicase-DNA-binding protein 1 homolog, CHD1 | 10.61 | 0.30 |  |
| PF3D7_1459000 | ATP-dependent RNA helicase DBP5 | 10.24 | 0.46 |  |
| PF3D7_1227100 | DNA helicase 60, DH60 | 6.55 | 0.41 | - |
| PF3D7_1007700 | AP2 domain transcription factor AP2-I | 4.65 | 0.08 | Invasion (Santos *et al.* 2017, Josling *et al.* 2020) |
| PF3D7_1107800 | AP2 domain transcription factor | 3.11 | 0.36 | Master regulator of parasite growth, chromatin structure and *var* gene expression (Subudhi *et al.* 2023) |
| PF3D7_0613800 | AP2 domain transcription factor | 2.43 | 0.28 | - |
| PF3D7_0420300 | AP2 domain transcription factor (ApiAP2) | 2.38 | 0.48 | - |
| PF3D7_0624600 | SNF2 helicase, ISWI | 2.09 | 0.001 | *var* gene expression (Bryant *et al.* 2020) |
| PF3D7_1239200 | AP2 domain transcription factor | 2.01 | 0.25 | - |

### PfMORC localizes to multigene families in subtelomeric regions

To determine the genome-wide occupancy of the *Pf*MORC chromatin-associated remodeling complex, we used chromatin immunoprecipitation followed by high-throughput sequencing (ChIP-seq). Using purified, crosslinked nuclear extracts, we immunoprecipitated *Pf*MORC from a 3xHA-tagged *Pf*MORC^HA-glmS^ parasite line (Singh *et al*. 2021) for ChIP-seq at the trophozoite stage (30hpi) and the schizont stage (40hpi) in biological duplicates. These timepoints represent the stages at which *Pf*MORC expression is highest (Singh *et al*. 2021). An independent ChIP-seq experiment in biological duplicate using anti-GFP and a *Pf*MORC^GFP^ parasite line (Singh *et al*. 2021) at the schizont stage was used to confirm our findings, demonstrating that the protein tags do not affect *Pf*MORC immunoprecipitation or genome-wide localization (**Supplemental Figure 2A,B**). As an additional control, we correlated one no-epitope (3D7 wild type), negative control sample using the same anti-HA antibody on unmodified parasite lines for immunoprecipitation (**Supplemental Figure 2A,B**; (Bonnell *et al*. 2023)), which resulted in an expected low correlation to the tagged samples. The biological ChIP-seq replicates showed high fold enrichment (Log_2_[IP/Input]) (**Supplemental Figure 2C**) and were highly correlated with each other within each timepoint (**Supplemental Figure 2D-F**).

We identified PfMORC localized to subtelomeric regions on all chromosomes across the *P. falciparum* genome, with additional occupancy at internal heterochromatic islands (**Figure 2A,B**). Within the subtelomeric regions, *Pf*MORC was bound upstream and within the gene bodies of many hypervariable multigene families (**Figure 2C**), including *var* genes (**Supplemental Figure 3**), *rif* genes (**Supplemental Figure 4**), and *stevor* genes (**Supplemental Figure 5**). Predicted binding sites (across both biological replicates) between the 30hpi and 40hpi timepoints showed a high degree of overlap, suggesting that *Pf*MORC binds many of the same regions throughout the later stages of asexual development when *Pf*MORC is highly expressed (**Figure 2D**). The proportion of *Pf*MORC-bound regions was similar across the 5′ upstream region of genes and the gene bodies throughout the genome, including subtelomeric regions (**Figure 2E**). As opposed to the binding of other proteins at the subtelomeric regions, such as the heterochromatin protein 1 (*Pf*HP1) (Flueck *et al*. 2010), *Pf*MORC occupancy is not widespread. Instead, it forms sharp peaks within, and adjacent to, HP1-bound regions (**Figure 2F**), suggesting a unique role for *Pf*MORC in heterochromatin condensation, boundary demarcation, and gene repression.

**Figure 2.**
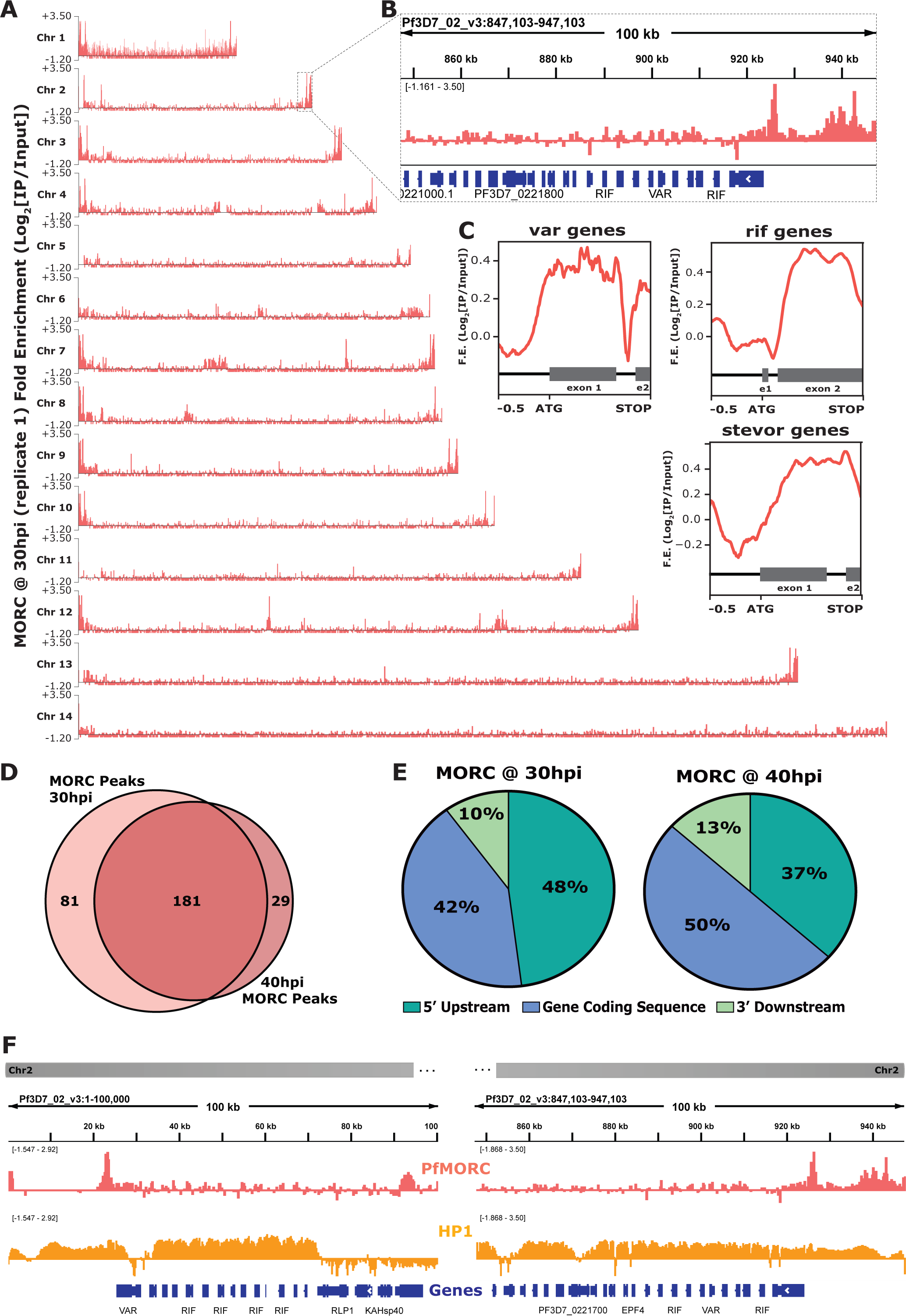
Genome-wide occupancy of *Pf*MORC reveals localization to hypervariable surface antigen genes at 30 h and 40 h. **(A)** Coverage tracks of *Pf*MORC across all 14 *P. falciparum* chromosomes. Plotted values are fold enrichment (Log2[IP/Input]) of a representative replicate at 30 h. **(B)** Zoom-in of the last 100kb region of chromosome two from Panel A. Gene annotations represented in blue bars (*P. falciparum* 3D7 strain, version 3, release 57; PlasmoDB.org). **(C)** Mean fold enrichment of *Pf*MORC occupancy across all var genes (top left), all rif genes (top right), and all stevor genes (bottom right), excluding pseudogenes. Graphical representation of exons to scale for each gene family annotated below enrichment plot in grey (e1=exon one; e2=exon two). **(D)** Quantitative Venn diagram comparing the number of MACS2 called peaks across each timepoint (light pink for 30 h; dark pink for 40 h). **(E)** Pie charts showing the type of genomic locations *Pf*MORC peaks overlap at both 30 h and 40 h. Pink slices are 5’ regions upstream of the ATG start site of genes, blue slices are coding sequences/gene bodies of genes, and green slices are 3’ regions downstream of the stop codon of genes. **(F)** Zoom-in of the first 100kb region (left) and the last 100kb region (right) of chromosome two. Plotted are the ChIP-seq fold enrichment of *Pf*MORC (top track; pink) and heterochromatin protein 1 (HP1; middle track; orange) with gene annotations (bottom track; blue bars; *P. falciparum* 3D7 strain, version 3, release 57; PlasmoDB.org).

We further defined *Pf*MORC putative gene targets as genes displaying peaks within 2-kb upstream of the ATG start codon or within gene bodies. For those peaks between gene targets in a head-to-head orientation, the closest gene was chosen. This resulted in 149 putative gene targets at 30hpi and 102 gene targets at 40hpi. A close examination of the 84 overlapping genes shows that many are *var* genes, rRNA genes, and genes encoding exported proteins (**Supplemental Table 4**), with GO terms related to cell adhesion, host‒pathogen interactions, and antigenic variation (**Supplemental Table 5**). The 65 uniquely bound genes at the 30hpi timepoint showed an enrichment of highly expressed tRNA and rRNAs genes, as well as conserved unknown genes, while those 18 unique to the 40hpi timepoint included a variety of late-stage expressed genes. Transcript abundance (Chappell *et al*. 2020) of the predicted *Pf*MORC gene targets at both the 30hpi and 40hpi timepoints form two major clusters: cluster one being genes expressed at the late ring/early trophozoite stage (10-24hpi) and cluster two at the late schizont stage (40-48hpi) (**Supplemental Figure 6**). This two-cluster gene target pattern of expression mirrors the biphasic pattern of expression by *Pf*MORC, suggesting that *Pf*MORC could have distinct functions, forming complexes with different sets of transcriptional regulators, at various times during asexual proliferation. As determined in other eukaryotic organisms, MORC family proteins do not generally bind DNA in a sequence-specific manner; it is, therefore likely that *Pf*MORC is recruited to these genome-wide regions by sequence-specific transcription factors, such as the ApiAP2 proteins identified in our proteomics results.

### Binding sites of PfMORC overlap with ApiAP2 proteins and epigenetic marks

*Pf*MORC has previously been found to interact with several ApiAP2 proteins (Hillier *et al*. 2019, Bryant *et al*. 2020, Singh *et al*. 2021), as does the *Toxoplasma* orthologue (Farhat *et al*. 2020, Srivastava *et al*. 2023, Antunes *et al*. 2024). We identified a clear overlap between genome-wide *Pf*MORC binding and putatively interacting ApiAP2 proteins using available ChIP-seq datasets. In addition to our protein-protein interaction results (**Table 1**), previous studies have also suggested that *Pf*MORC interacts with a broad array of ApiAP2 TFs, such as *Pf*AP2-G5, *Pf*AP2-O5, *Pf*AP2-I, PF3D7_1107800, PF3D7_0613800, PF3D7_0420300, and PF3D7_1239200 (Hillier *et al*. 2019, Bryant *et al*. 2020, Subudhi *et al*. 2023). Therefore, we compared binding sites between interacting ApiAP2s and *Pf*MORC using available ChIP-seq data on *Pf*AP2-G5, *Pf*AP2-O5, *Pf*AP2-I, PF3D7_1107800, PF3D7_0613800, PF3D7_1239200 (Josling *et al*. 2020, Shang *et al*. 2021, Shang *et al*. 2022). Interestingly, there is a degree of overlap between the binding sites of all six ApiAP2 TFs and *Pf*MORC, suggesting that *Pf*MORC and these ApiAP2 TFs may cooperate in the regulation of gene expression at these loci (**Figure 3A-B**; **Supplemental Figure 7**). However, the available data cannot differentiate whether all these factors are in one complex together, form multiple smaller heterologous complexes, or are components of separate complexes in individual cells. DNA motif enrichment analysis (Bailey 2021) identified several unique and significant DNA motifs at both the 30hpi and 40hpi timepoints, which suggests that more than one sequence-specific transcription factor may be responsible for recruiting *Pf*MORC to specific genomic regions (**Supplemental Figure 8**). The common motifs identified across replicates and timepoints are RGTGCAW or TGCACACA, both of which are similar or identical to the *in vitro* and/or *in vivo* DNA-binding motif of *Pf*AP2-I (RGTGCAW) or PF3D7_0420300 (TGCACACA), respectively, suggesting that these ApiAP2 factors may play major roles in *Pf*MORC recruitment (**Supplemental Figure 8**).

**Figure 3.**
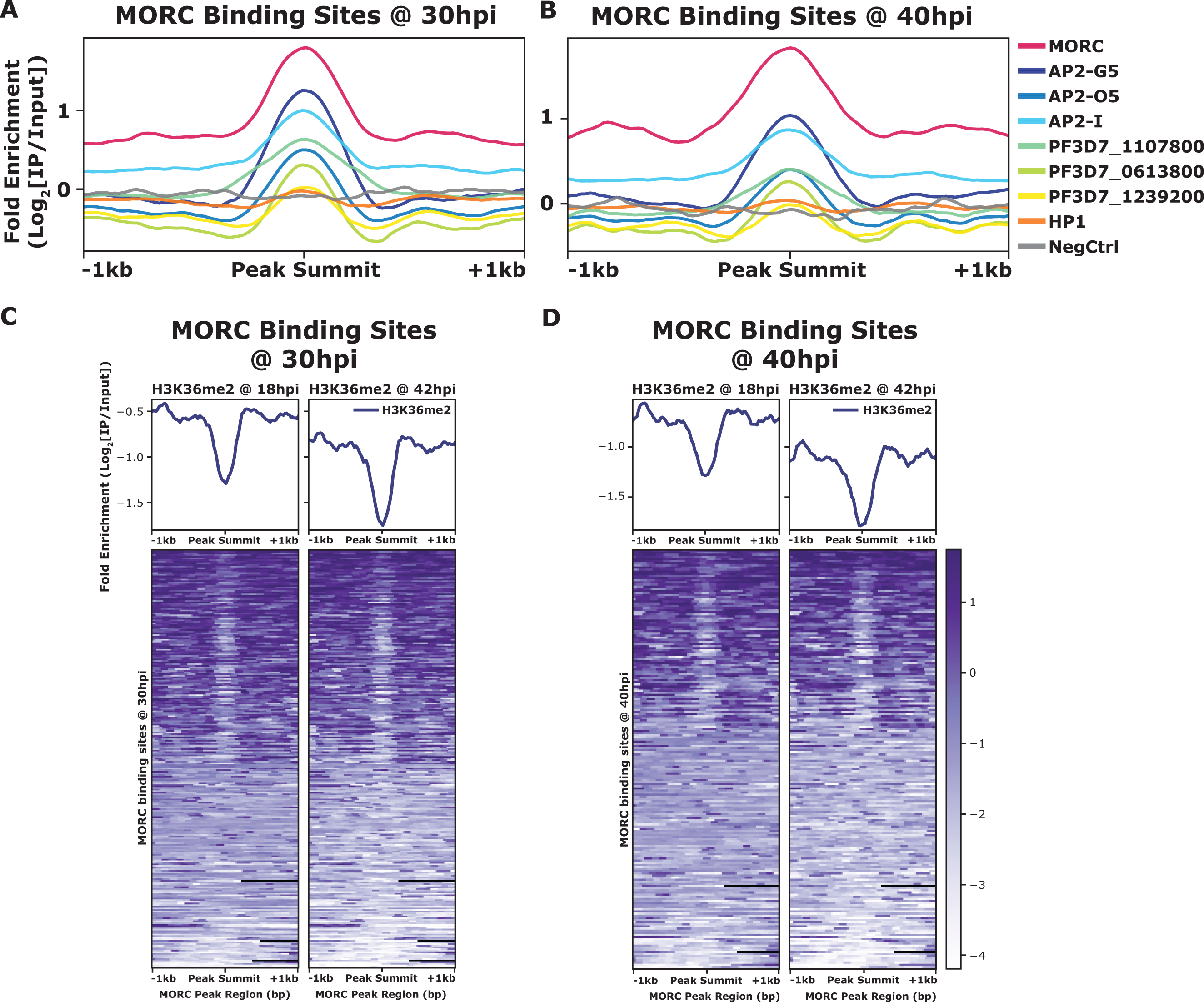
**(A)** Mean fold enrichment (Log2[IP/Input]) of *Pf*MORC, six associated factors (AP2-G5, AP2-O5, AP2-I, PF3D7_1107800, PF3D7_0613800, and PF3D7_1239200), HP1, and a negative no-epitope control across *Pf*MORC binding sites at the 30 h timepoint. **(B)** Mean fold enrichment (Log2[IP/Input]) of *Pf*MORC, six associated factors (AP2-G5, AP2-O5, AP2-I, PF3D7_1107800, PF3D7_0613800, and PF3D7_1239200), HP1, and a negative no-epitope control across *Pf*MORC binding sites at the 40 h timepoint. **(C)** Mean fold enrichment (Log2[IP/Input]) and heatmap of two H3K36me2 epigenetic mark timepoints across *Pf*MORC binding sites at 30 h. **(D)** Mean fold enrichment (Log2[IP/Input]) and heatmap of two H3K36me2 epigenetic mark timepoints across *Pf*MORC binding sites at 40 h.

In addition to overlapping occupancy with ApiAP2 TFs, we found that *Pf*MORC co-localizes with the depletion of H3K36me2 (**Figure 3C, D**), which is demarcated by the SET2 methyltransferase (Jiang *et al*. 2013), both at 30hpi and 40hpi. No other significant co-localization was found between *Pf*MORC and other epigenetic marks (H2A.Z, H3K9ac, H3K4me3, H3K27ac, H3K18ac, H3K9me3, H3K36me2/3, H4K20me3, and H3K4me1) (**Supplemental Figure 9**), suggesting it has a unique binding preference not shared with other heterochromatin markers. Therefore, it is likely that *Pf*MORC co-localizes with other, as yet uncharacterized, epigenetic marks. In summary, *Pf*MORC was found to be recruited to 5’- untranslated regions (UTRs), gene body regions, and subtelomeric regions of repressed, multigene families, and overlaps with other known ApiAP2 binding sites and DNA motifs.

### Depletion of PfMORC results in upregulation of late-stage genes associated with invasion

*Pf*MORC association with chromatin remodelers has been shown (Bryant *et al*. 2020), but how *Pf*MORC regulates gene expression in the asexual stage has not been evaluated. In *T. gondii*, the *Tg*MORC:*Tg*ApiAP2 complex acts as a transcriptional repressor of sexual commitment (Farhat *et al*. 2020, Srivastava *et al*. 2023, Antunes *et al*. 2024). Here, we found that *Pf*MORC co-immunoprecipitates with several chromatin remodeling proteins and many ApiAP2 transcription factors. Furthermore, our ChIP-seq data revealed that *Pf*MORC is located at subtelomeric regions of the genome. Based on this evidence, we hypothesized that PfMORC may regulate transcriptional changes during blood stage development of the parasite. To knock down the expression of *Pf*MORC (*Pf*MORC-KD), sorbitol-synchronized MORC^HA-glmS^ parasites (22-24hpi) were subjected to 2.5 mM glucosamine (GlcN) treatment for little over 48 hours when they reached the trophozoite stage (32hpi ± 3hpi), at which point parasites were harvested for RNA isolation for transcriptomic analysis. In parallel, another flask with *Pf*MORC^HA-glmS^ parasites was set up without GlcN and used as control for RNA-seq comparison. We previously demonstrated that treatment with 2.5 mM GlcN results in a 50% knockdown of *Pf*MORC protein but does not cause any growth delay; using >2.5 mM GlcN caused a measurable slow growth and reduced parasitemia (Singh *et al*. 2021). Three biological replicates with and without 2.5 mM GlcN were collected for knockdown transcriptomics to ensure reproducibility.

For each *Pf*MORC RNA-seq sample, gene counts were used to identify the differentially expressed genes (DEGs). Significant threshold parameters were assigned to a p value <0.05, yielding a total of 2558 DEGs (**Supplemental Table 6**). Applying a log_2_-fold change cut-off from >1 to <-1 and filtering out pseudogenes reduced this number to 163 DEGs. Among these, 60 genes display more abundant transcripts, whereas 103 genes were reduced relative to control parasites grown without GlcN **(Figure 4A)**. Pathway and functional enrichment analysis yielded several genes from apical organelles. GO analysis revealed gene clusters enriched with molecular function (p-adj = 0.0006) involved in the movement into the host environment and entry into the host, and molecular function of protein binding (p-adj = 0.009) (**Figure 4B**, **Supplemental Table 7)**. More specifically, upregulated genes implicated in the invasion of merozoites were found to be expressed prematurely upon *P****f***MORC KD; these include several rhoptry-associated genes, notably *Pf*RON2 (PF3D7_1452000) and *Pf*RON3 (PF3D7_1252100). Both *Pf*RON2 and *Pf*RON3 are part of a micronemal complex at the erythrocyte membrane where *Pf*RON2 anchors *Pf*AMA1 to facilitate merozoite invasion (Srinivasan *et al*. 2013). In addition, *Pf*SUB3 (PF3D7_0507200), *Pf*SERA5 (PF3D7_0207600) and *Pf*DPAP3 (PF3D7_0404700), all of which are critical for schizont rupture (Yeoh *et al*. 2007, Arastu-Kapur *et al*. 2008), were among the upregulated DEGs (**Figure 4C**). In general, we found that depletion of *Pf*MORC leads to upregulation of invasion-related genes, which suggests that *Pf*MORC has an additional function in the regulation of genes specifically associated with red blood cell invasion.

**Figure 4.**
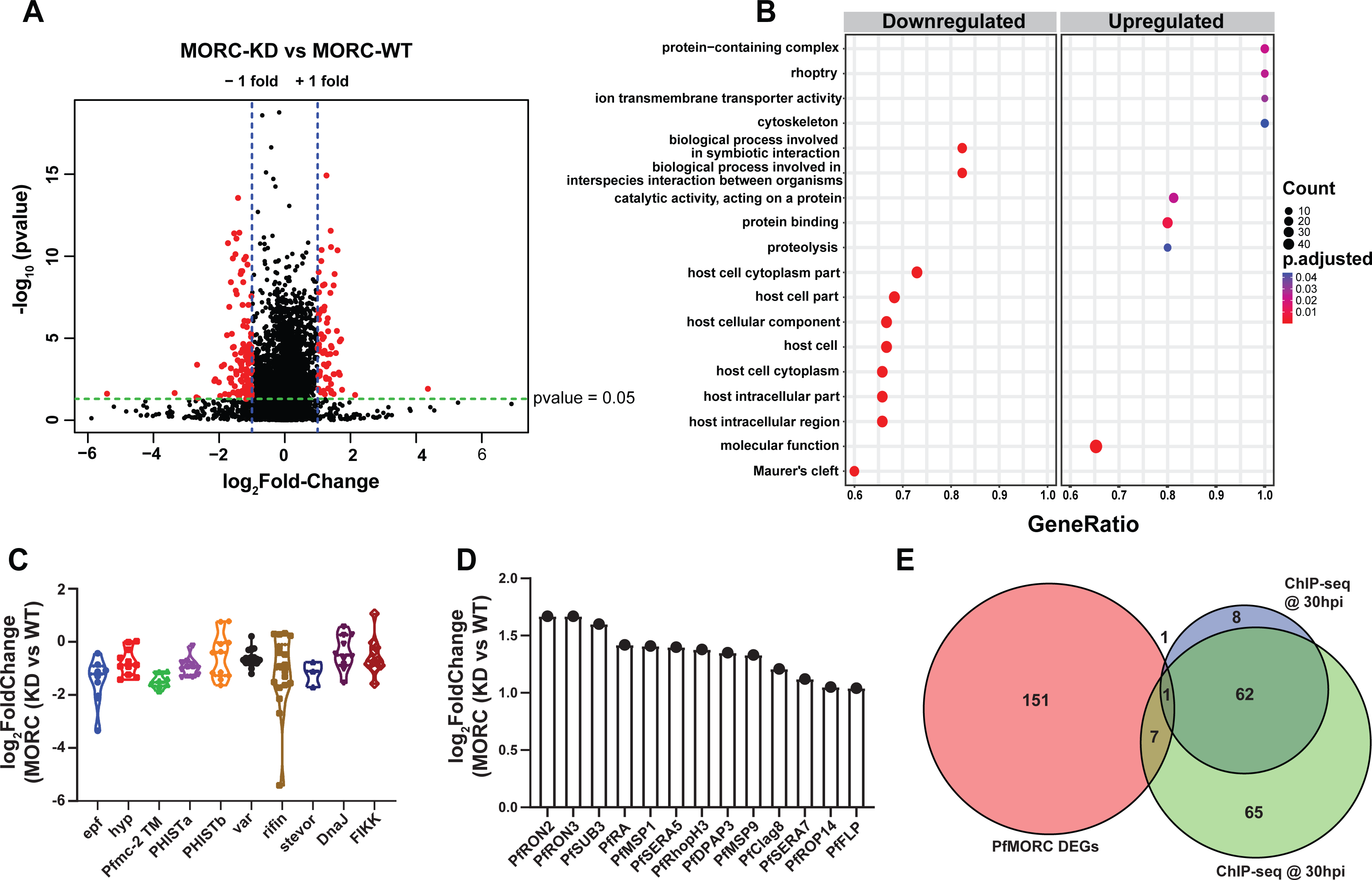
Transcriptome analysis of *Pf*MORC knockdown revealed differential gene expression. **(A)** Volcano plot displaying the differential gene expression in *Pf*MORC-KD compared to the *Pf*MORC-WT phenotype. Tightly synchronized PfMORC^HA-glmS^ parasites (32 hpi ± 3 h) were split into two populations, one of which was treated with 2.5 mM GlcN to obtain the *Pf*MORC knockdown phenotype and the other was not treated with GlcN to obtain wild-type phenotype. Total RNA-seq was performed, and significant threshold parameters for DEGs were assigned to a *p-value* < 0.05 and -log2 fold change > 1 from three biological replicates. **(B)** Scatter plot shows upregulated and downregulated DEGs which were further categorized for pathway and functional enrichment analysis using the KEGG database (p-adjusted value < 0.05). The circle size at the vertical axis represents the number of genes in the enriched pathways and the horizontal axis represents gene richness as a ratio of DEGs in the pathways to the total genes in a specific pathway. **(C)** The violin plot of log_2_ fold change of genes belonging to the multigene family is constructed from *Pf*MORC-KD vs *Pf*MORC-WT, which shows DEGs of multigene family proteins upon *Pf*MORC knockdown. **(D)** The bar plot illustrates the upregulated DEGs of apical organelle origin in *Pf*MORC-KD parasites involved in host cell invasion. **(E)** Venn diagram showing the comparison between genes obtained from ChIP-seq data and DEGs obtained from RNA-seq data. Both 30hpi and 40 hpi time points were taken for comparison and showed high overlap with each other but there was no overlap with RNA-seq data.

### PfMORC-depleted parasites downregulate hypervariable gene families

Among the genes with reduced mRNA abundance, DEGs linked to cytoadherence, antigenic variation and interaction with host (**Figure 4B and D**) where over-represented in the GO analysis. Many of the genes enriched in the downregulated group belong to the clonally variant *var* multigene family that represents 60 members encoding the *P. falciparum* erythrocyte membrane proteins (*Pf*EMP1), which upon switching expression, aid in pathogenesis and immune evasion (Guizetti and Scherf 2013). Furthermore, a cluster of genes encoding putative exported proteins were also enriched including members of the exported protein family (EPFs), *Plasmodium* exported protein (hyp), and *Plasmodium* exported protein (PHISTa/b). Other significantly overrepresented downregulated genes belonged to serine/threonine protein kinases, the FIKK family (Ward *et al*. 2004), most of which are exported to the RBC, and Maurer’s clefts two transmembrane proteins (Pfmc-2TM) (**Figure 4D**). Notably, expressed proteins are conserved across the *Plasmodium* family and remain confined to the hypervariable subtelomeric region of *P. falciparum* chromosomes (Sargeant *et al*. 2006).

### Comparison of ChIP-seq gene targets and DEGs by RNA-seq

To identify whether genes found to be dysregulated after PfMORC knockdown are associated with the genome-wide occupancy of *Pf*MORC, we correlated the gene targets identified by ChIP-seq with the DEGs determined by RNA-seq. We identified a total of 135 gene targets from the 30hpi ChIP-seq timepoint, 72 gene targets from the 40hpi ChIP-seq timepoint, and 163 DEGs by RNA-seq. The low correlation between the ChIP-seq gene targets and the RNA-seq DEGs suggest that *Pf*MORC genome-wide occupancy is more likely involved in chromatin structure, rather than specific regulation of gene targets (**Figure 4E**). Likely, *Pf*MORC localizes to these sites to aid in chromatin condensation, as shown in other eukaryotic systems (Zhang *et al*. 2019, Zhong *et al*. 2023).

## Discussion

Periodic gene expression during the asexual blood stage directly corresponds to the timing in which gene products are needed (Bozdech *et al*. 2003) and shows oscillation patterns associated with circadian rhythms (Smith *et al*. 2020, Subudhi *et al*. 2020). Transcription factors are critical regulators of this dynamic pattern in concert with epigenetic regulators and genome-wide changes to chromatin structure (Painter *et al*. 2011, Toenhake *et al*. 2018, Jeninga *et al*. 2019, Hollin *et al*. 2021). The ApiAP2 family members display unique binding preferences in the genome (Campbell *et al*. 2010), undergo stage-specific expression (Painter *et al*. 2011), and have been identified to regulate virtually all stages of development across multiple *Plasmodium* species (Jeninga *et al*. 2019). To date, ApiAP2 proteins have been reported in transcriptional silencing of clonally variant genes, regulation of invasion genes, and sexual commitment, acting through interaction with other epigenetic factors, such as heterochromatin protein 1 [*Pf*HP1 (Flueck *et al*. 2010, Brancucci *et al*. 2014, Fraschka *et al*. 2018)], bromodomain protein 1 [*Pf*BDP1 (Santos *et al*. 2017, Josling *et al*. 2020, Quinn *et al*. 2022)], or general control non-depressible 5 [*Pf*GCN5 (Miao *et al*. 2021)]. Despite the known DNA-binding sites of many ApiAP2 proteins (Flueck *et al*. 2010, Santos *et al*. 2017, Sierra-Miranda *et al*. 2017, Josling *et al*. 2020, Carrington *et al*. 2021, Shang *et al*. 2021, Singh *et al*. 2021, Russell *et al*. 2022, Shang *et al*. 2022, Bonnell *et al*. 2023), limited information is available for other accessory proteins.

Recently, the *T. gondii* orthologue *Tg*MORC was identified in a complex with HDAC1, AP2XII-2, and AP2XII-1:AP2XI-2and orchestrates epigenetic rewiring of sexual gene transcription (Farhat *et al*. 2020, Srivastava *et al*. 2023, Antunes *et al*. 2024). In *P. falciparum*, *Pf*MORC has been copurified with several ApiAP2 proteins, as shown by different independent studies (Hillier *et al*. 2019, Bryant *et al*. 2020, Singh *et al*. 2021).

In this study, we used an integrated multi-omics approach to explore the function of *Pf*MORC during asexual blood stage development. Using immunoaffinity-based purification, we identified a number of nuclear proteins that interact with *Pf*MORC. More specifically, *Pf*MORC was associated with *Pf*AP2-G5, which is essential for gametocytogenesis (Shang *et al*. 2021), *Pf*AP2-I, which is required for the expression of many invasion-related genes (Santos *et al*. 2017), and with other ApiAP2 TFs of unknown function (*Pf*AP2-O5, PF3D7_1107800, PF3D7_0613800, PF3D7_0420300, and PF3D7_1239200). The identification of *Pf*ISW1 and *Pf*CHD1 in association with *Pf*MORC strengthens the link between *Pf*MORC and chromatin remodeling. This finding suggests that *Pf*MORC may participate in regulating chromatin structure and gene expression during the IDC, notably the specific regulation of *var* genes. We note that some nuclear proteins were not identified in this study despite their documented interaction with *Pf*MORC in other studies (Hillier *et al*. 2019, Bryant *et al*. 2020, Singh *et al*. 2021, Subudhi *et al*. 2023). This may be due to differences in experimental conditions between the various studies, and to the low abundance of the proteins. Despite this, our identification of several nuclear proteins in complex with *Pf*MORC provides insights into potential interactions and regulatory mechanisms underlying gene regulation and chromatin remodeling in *P. falciparum*; this may have implications for developing new strategies to combat malaria.

In other eukaryotes, MORC proteins function in gene repression and chromatin compaction at heterochromatic regions (Koch *et al*. 2017). Therefore, to determine if *Pf*MORC localizes to heterochromatic regions across the *P. falciparum* genome, we performed ChIP-seq assays with MORC^HA-glmS^ parasites during peak PfMORC expression (trophozoite and schizont stages). In general, *Pf*MORC occupied regions 5’–upstream of the ATG start site or was bound within coding region, irrespective of the developmental stage. Most importantly, *Pf*MORC peaks were reproducibly detected in subtelomeric regions containing hypervariable multigene families, including the *var* genes, consistent with the findings that *Pf*MORC localized to *var* gene promoters as reported using dCas9 (Bryant *et al*. 2020). The specific binding pattern of *Pf*MORC near or within *Pf*HP1-bound regions suggests two related critical functions: heterochromatin condensation and gene repression. It is possible that *Pf*MORC changes the nucleosome landscape either by direct association or by binding to different chromatin remodelers. Recent work in *Arabidopsis* has further confirmed the importance of *At*MORC paralogs in gene regulation by chromatin compaction (Zhong *et al*. 2023), which resembles the function of *Pf*MORC in *P. falciparum*. Interestingly, in *T. gondii*, the major function of *Tg*MORC was in the repression of sex-determination genes (Farhat *et al*. 2020), suggesting MORC family proteins have the capacity to perform diverse functions across eukaryotic organisms. Future studies should determine if *Pf*MORC depletion similarly influences the rate of sexual commitment to *P. falciparum* gametocytogenesis, which is known to be regulated by epigenetic mechanisms. Role of *Pf*HP1 has been shown in regulating sexual commitment by repressing *Pf*AP2-G and virulence genes expression (Brancucci *et al*. 2014), whereas gametocyte development 1 (*Pf*GDV1) displaces *Pf*HP1 itself and induces asexual developing parasites to sexual differentiation (Filarsky *et al*. 2018).

We also compared *Pf*MORC occupancy with available ChIP-seq data for the interacting ApiAP2 proteins. Interestingly, our analysis revealed a significant overlap between the binding sites of all ChIP-ed ApiAP2 proteins with *Pf*MORC, suggesting a cooperative role in gene regulation. We also identified enriched motifs similar to those bound by our ApiAP2 proteins of interest, further suggesting the functional cooperation between *Pf*MORC and ApiAP2 proteins. Of note, only one of the seven ApiAP2 of interest in our study has not been ChIP-ed to date (PF3D7_0420300) (Shang *et al*. 2022). However, we found an enrichment of the DNA motif (TGCACACA) at *Pf*MORC-bound sites. This motif is bound *in vitro* by PF3D7_0420300 (Bonnell *et al*. 2023), suggesting that this ApiAP2 protein may localize to these regions. Overall, since these seven ApiAP2 proteins are expressed at distinct time points during the *P. falciparum* cycle (Bozdech *et al*. 2003) and regulate different sets of genes (Santos *et al*. 2017, Josling *et al*. 2020, Shang *et al*. 2021, Shang *et al*. 2022, Subudhi *et al*. 2023), we believe this indicates a variety of functions for *Pf*MORC at different stages. In addition, a recent study in *Arabidopsis* reported that *At*MORC-mediated regulation of transcription may be due to both direct chromatin interactions and indirect association via sequence-specific transcription factors (Zhong *et al*. 2023) consistent with a complex landscape of chromatin remodeling by MORC proteins. We also interrogated the co-localization of *Pf*MORC and numerous epigenetic profiles (H2A.Z, H3K9ac, H3K4me3, H3K27ac, H3K18ac, H3K9me3, H3K36me2/3, H4K20me3, and H3K4me1), with a specific focus on H3K36me2, since this histone modification has been shown to act as a global repressive effector in *P. falciparum* (Karmodiya *et al*. 2015) and other organisms (Strahl *et al*. 2002, Wagner and Carpenter 2012). Interestingly, we did not find a strong co-localization with any of these epigenetic marks, suggesting a unique role of the epigenetic landscape on *Pf*MORC global occupancy. Therefore, the functional association between *Pf*MORC and a specific, unknown epigenetic mark remains to be determined.

Before this work, there was no direct evidence correlating *Pf*MORC-mediated transcriptional changes during the IDC of *P. falciparum*. Therefore, we investigated the differentially expressed genes (DEGs) in *Pf*MORC knockdown parasites and found two distinct subsets of enriched genes. The upregulated DEGs were enriched with genes related to invasion, while the downregulated DEGs belong to hypervariable genes associated with parasite virulence. It suggests that *Pf*MORC has very dynamic function across *Plasmodium* asexual cycle as we also identified *Pf*AP2-I in CoIPed proteins, which is shown to regulate invasive gene transcription (Santos *et al*. 2017). Overall, our data with *Pf*MORC-KD revealed a change in the expression of both antigenically variable and invasion-related genes. Expression of clonally variant genes occurs in a different but tightly regulated manner in IDC (Scherf *et al*. 1998, Scherf *et al*. 2008), which, in the light of our data, may be regulated by *Pf*MORC occupancy. Data regarding changes in *var* gene expression are difficult to interpret, as the KD experiments were performed on a parasite that had not been recloned and may therefore express more than one *var* gene. Single-cell experiments would be needed to clarify the effect of *Pf*MORC KD on *var* gene regulation. We were surprised by the small number of DEGs detected upon knockdown of *Pf*MORC, which we believe may be due to incomplete knockdown.

We previously showed that parasites in which *Pf*MORC is knocked-down display reduced sensitivity to melatonin (Singh et al. 2021). This prompted us to investigate if overall gene expression in *Pf*MORC KD parasites is affected by melatonin treatment. We detected only very slight changes **(Supplemental Figure 10A-D)**, suggesting that melatonin does not exert its effect on gene expression through *Pf*MORC, or at least that the latter protein plays a minor role in the process.

Overall, this study shows that *Pf*MORC interacts with different ApiAP2 TFs, in line with previous studies (Hillier *et al*. 2019, Bryant *et al*. 2020, Singh *et al*. 2021, Subudhi *et al*. 2023) and a recently published parallel study (Chahine *et al*. 2023). Furthermore, we found that *Pf*MORC is localized at sub-telomeric regions and contains significant overlap with the bindingsites of several ApiAP2 transcription factors. Our results support a role for *Pf*MORC in the regulation of hypervariable genes that are essential for *Plasmodium* virulence and rhoptry genes critical to red blood cell invasion. Collectively, our data identify an important role for *Pf*MORC in chromatin organization, as well as in the epigenetic regulation of gene expression through regulatory complexes with an array of transcription factors, making it an attractive drug target.

## Materials and Methods

### Plasmodium falciparum culture

The *P. falciparum* strains were cultured at 37°C in RPMI 1640 medium supplemented with 0.5% Albumax II (Gibco) (Trager and Jensen 1976). Parasites were grown under a 5% CO_2_, 5% O_2_, and 90% N_2_ atmosphere. Cultures were synchronized with 5% sorbitol (Lambros and Vanderberg 1979).

### Coimmunoprecipitation and mass spectrometry

Infected erythrocytes at the trophozoite stage were collected from culture and washed twice in 1x phosphate-buffered saline (PBS). The culture pellet was suspended in PBS containing 0.05% (w/v) saponin to lyse the erythrocyte membrane and centrifuged at 8,000 × g for 10 min. The supernatant was discarded, and the parasite pellet was washed 3 times in cold PBS. To perform the co-immunoprecipitation, we followed the manufacturer protocol (ChromoTek, gta-20). Samples were lysed in modified RIPA buffer (50 mM Tris, pH 7.5, 150 mM NaCl, 0.5% sodium deoxycholate, 1% Nonidet P-40, 10 µg/ml aprotinin, 10 µg/ml leupeptin, 10 µg/ml, 1 mM phenylmethylsulfonyl fluoride, benzamidine) for 30 min on ice. The lysate was precleared with 50 µl of protein A/G-Agarose beads at 4°C for 1 h and clarified by centrifugation at 10,000 × g for 10 min. The precleared lysate was incubated overnigh with an anti-GFP-Trap-A beads (ChromoTek, gta-20) antibody. The magnetic beads were then pelleted using a magnet (Invitrogen), and the beads were washed extensively using wash buffer (50 mM Tris, pH 7.5, 150 mM NaCl, 0.5% sodium deoxycholate, 1% Nonidet P-40) to minimize the detection of non-specific proteins. To elute the immunoprecipitated proteins, the magnetic beads were resuspended in 2X SDS loading buffer and resolved by SDS‒PAGE. Following SDS‒PAGE, the whole gel band for each sample was excised and further analysed by mass spectrometry. We used a service provider (CEFAP core-facility de Espectometria de Massa) to analyse GFP-coimmunoprecipitated proteins.

### In-gel digestion and peptide desalting

Protein bands from polyacrylamide gels were cut into pieces (approximately 1 mm^3^), transferred to a clean 1.5 ml low binding tube and washed with washing solution (40% acetonitrile, 50 mM ammonium bicarbonate) until the bands were completely distained, and dehydrated with ACN 100% for 5 min followed by vacuum centrifugation. Proteins were then reduced with 10 mM dithiothreitol (DTT) in 50 mM ammonium bicarbonate and incubated for 45 minutes at 56°C. Protein alkylation was performed with 55 mM iodoacetamide (IAA) in 50 mM ammonium bicarbonate and incubated for 30 minutes at room temperature. Proteins were digested into peptides by trypsin overnight reaction at 37°C. The trypsin reaction was stopped with 10% TFA (1% TFA final concentration). The supernatant was collected into a new tube. Extraction buffer (40% ACN/0.1% TFA) was added to the gel pieces and incubated for 15 minutes on a thermomixer at room temperature. The supernatant was transferred to the same microtube. The peptide extraction was performed twice and then dried in a vacuum centrifuge. Peptides were resuspended in 0.1% TFA for desalting.

### Nano LC‒MS/MS analysis

The LC‒MS/MS system employed was an Easy-nano LC 1200 system (Thermo Fisher Scientific Corp) coupled to an Orbitrap Fusion Lumos mass spectrometer equipped with a nanospray source (Thermo Fisher Scientific Corp). Samples were loaded onto a trapping column (Acclaim PepMap 0.075 mm, 2 cm, C18, 3 μm, 100 A; Thermo Fisher Scientific Corp.) in line with a nano-LC column (Acclaim PepMap RSLC 0.050 mm, 15 cm, C18, 2 μm, 100 A; Thermo Fisher Scientific Corp.). The gradient was 5-28% solvent B (A 0.1% FA; B 90% ACN, 0.1% FA) for 25 minutes, 28-40% B for 3 minutes, 40-95% B for 2 minutes and 95% B for 12 minutes at 300 nL/min. Orbitrap Fusion Lumos mass spectrometer operated in positive mode. The full MS scan had an automatic gain control (AGC) of 5 x 105 ions and a maximum filling time of 50 ms. Each MS scan was acquired at 120K full width half maximum (FWHM) high resolution in the Orbitrap with a mass range of m/z 400-1600 Da. High-resolution dissociation (HCD) with a normalized collision energy set at 30 was used for fragmentation. The resulting MS/MS fragment ions were detected in the Orbitrap mass analyser at a resolution of 30,000. An AGC of 5 x 104 ions and a maximum injection time of 54 ms were used. All raw data were accessed in Xcalibur software (Thermo Scientific).

### Database searches and bioinformatics analyses

Raw files were imported into MaxQuant version 1.6.17.0 for protein identification and quantification. For protein identification in MaxQuant, the database search engine Andromeda was used against the UniProt *P. falciparum* 3D7 strain (March 2021, 5388 entries release). The following parameters were used: carbamidomethylation of cysteine (57.021464 Da) as a fixed modification, oxidation of methionine (15.994915 Da) and N-terminal acetylation protein (42.010565 Da) were selected as variable modifications. Enzyme specificity was set to full trypsin with a maximum of two missed cleavages. The minimum peptide length was set to 7 amino acids. For label-free quantification, the “match between runs” feature in MaxQuant was used, which is able to identify the transfer between samples based on the retention time and accurate mass, with a 0.7-minute match time window and 20-minute alignment time window. Label-free quantification was performed using MaxQuant software with the “match between run” and iBAQ features activated. Protein LFQ and iBAQ ratios were calculated for the two conditions, and the protein IDs were divided accordingly. The mass spectrometry proteomics data have been deposited to the ProteomeXchange Consortium via the PRIDE (Perez-Riverol *et al*. 2022) partner repository with the dataset identifier PXD036092. PlasmoDB database was used to perform GO analysis.

### Total RNA extraction and RNAseq

Infected RBCs containing tightly synchronized trophozoite stage parasites (32 hpi ± 3 h) were harvested and resuspended in TRIzol (Thermo Fisher Scientific). Total RNA was extracted following the manufacturer’s protocol, and an RNA cleanup kit (Qiagen) was used to achieve high-purity RNA. The isolated RNA was quantified using a NanoDrop ND-1000 UV/Vis spectrophotometer, and RNA quality was determined using an RNA ScreenTape System (Agilent 2200 TapeStation). Total RNA (10000 ng) from each sample was stabilized in RNAStable (Biomatrica) and sent to the Micromon Genomics facility at Monash University for next-generation sequencing.

RNA samples were prepared using the MGITech RNA Directional Library Prep Kit V2 (Item No. 1000006385), as per the manufacturer’s instructions, with the following parameters: input RNA: 50 ng, fragmentation: target of 200 bp-400 bp, 87 °C for 6 mins, adapter clean-up: 200 bp-400 bp and library amplification cycles: 16. Libraries were assessed for quantity using the Invitrogen Qubit and dsDNA HS chemistry (Item No. Q33230) and quality/quality using the Agilent Fragment Analyser 5200 with the HS NGS Fragment Kit (Item No. DNF-473-0500). The libraries were pooled in equimolar concentrations and sequenced using an MGITech DNBSEQ-G400RS sequencing instrument with High-Throughput Sequencing Set (FCL PE100, Item No. 1000016950) chemistry.

The quality of the RNA-seq libraries was evaluated using the FastQC tool. Next, we used Salmon (V1.9.0) (Patro *et al*. 2017) quant with default arguments to quantify the expression of all transcripts in the PlasmoDB release-58 Pfalciparum3D7 genome. The transcript expression was summarized to gene-level expression with tximport 1.22.0 (Soneson *et al*. 2015). Finally, the gene counts were used to detect differentially expressed genes (DEGs) with DESeq2 (1.34.0) (Love *et al*. 2014). Furthermore, only genes with an >1 log_2_ fold change and adjusted *p-value* <0.1 were considered significant for further analysis.

### Chromatin immunoprecipitation followed by high-throughput sequencing (ChIP-seq)

PfMORC ChIP-seq was performed similarly to previously published ChIP-seq experiments in *Plasmodium falciparum* (Josling *et al*. 2020, Singh *et al*. 2021, Russell *et al*. 2022). Five samples in total were collected: biological duplicates using the PfMORC^HA-glmS^ parasite line at the trophozoite stage [30 hpi], biological duplicates using the PfMORC^HA^ parasite line at the schizont stage [40 hpi], and a single replicate using the PfMORC^GFP^ parasite line at the schizont stage [40 hpi]. In brief, the protocol includes five steps: (1) Treated with 1% formaldehyde to crosslink the suspended PfMORC^HA-glmS^ (or PfMORC^GFP^) parasite cultures (at least 10^8^ trophozoite- or schizont-stage parasites synchronized with 10% sorbitol more than one cycle prior) at 37°C for 10 minutes; (2) Collected parasite nuclei using prechilled glass Dounce homogenizer for 100 strokes per 10^9^ trophozoites/schizonts; (3) Lysed parasite nuclei and mechanically sheared chromatin until sufficiently sheared using Covaris Focus-Ultrasonicator M220 (5% duty cycle, 75 W peak incident power, 200 cycles per burst, 7°C, for 5 minutes). (4) We collected 10% of each sample for the non-immunoprecipitated “input” control and then immunoprecipitated the remaining 90% of each sample. The remaining 90% of each sample was immunoprecipitated with 1:1000 anti-HA antibody (0.5 mg/mL Roche Rat Anti-HA High Affinity [11867423001]) or 1:1000 anti-GFP antibody (0.1 mg/mL Abcam Rabbit Anti-GFP [Ab290]) overnight at 4°C with rotation. (5) DNA was purified after reverse crosslinking using a MinElute column (Qiagen) as directed and quantified by a Qubit fluorometer (Invitrogen).

### ChIP-seq library prep for Illumina sequencing

The PfMORC ChIP-seq libraries were performed similarly to previously published ChIP-seq experiments in *Plasmodium falciparum* (Josling *et al*. 2020, Singh *et al*. 2021, Russell *et al*. 2022). DNA sequencing libraries were prepared for high-throughput Illumina sequencing on the NextSeq 2000 with the 150 x 150 single-end mode. The library prep underwent 12 rounds of amplification using KAPA HiFi polymerase. Completed libraries were quantified using the Qubit fluorometer (Invitrogen) for high-sensitivity DNA and library sequence length by the Agilent TapeStation 4150.

### ChIP-seq data analysis and peak calling

The MORC ChIP-seq datasets were analysed similar to previously published ChIP-seq experiments in *Plasmodium falciparum* **(Josling *et al*. 2020, Singh *et al*. 2021, Russell *et al*. 2022)**. Raw sequencing reads were trimmed (Trimmomatic v0.32.3) to remove Illumina adaptor sequences and low-quality reads below 30 Phred (SLIDINGWINDOW: 4:30). FastQC (v0.11.9) was used to check the quality after trimming. Processed reads were then mapped to the *P. falciparum* genome (release 57) using BWA-MEM (v0.7.17.2) simple Illumina mode with multiple mapped reads filtered out (MAPQ=1). Once the sequences were mapped, MACS2 (v2.2.7.1) was used to call peaks with each biological replicate and its paired input sample using a standard significance cut-off (q-value=0.01). Using BedTools Multiple Intersect (v2.29.2), the narrow peak output file for each replicate was overlapped to identify the significant peaks in the overlap between both ChIP-seq biological replicates. The overlapping regions were then used to identify an enriched DNA motif (STREME Meme Suite: (Bailey 2021), and putative gene targets of PfMORC were defined as genes with peaks within 2 kb upstream of the gene target ATG start codon or peaks within gene bodies. In a situation with any peaks between gene targets in a head-to-head orientation, the closest gene was chosen.

## Data availability

The mass spectrometry proteomics data have been deposited to the ProteomeXchange Consortium via the PRIDE partner repository with the dataset identifier PXD044256. All ChIP-seq samples were submitted to GEO, and a GEO submission ID (GSE239393) was obtained for this section. Additionally, submit all the relevant ChIP-seq files to Zenodo, since UCSC browser doesn’t easily work from *Plasmodium falciparum* data. All RNA-seq samples were submitted to GEO, and a GEO submission ID (GSE241313) was obtained for this section.

## Author Contributions

MKS, CRSG, and ML designed the experiments. MKS, VAB, VFS and MSM performed the experiments, and data were analyzed by MKS, VAB, ITS, GP, ML and CRSG. The primary manuscript was written by MKS and VAB and proofread by CD, ML and CRSG.

## Acknowledgments

This work was supported by grants from Fundação de Amparo a Pesquisa de São Paulo (FAPESP) to C.R.S.G. (2017/08684-7 and 2018/07177-7) and to M.K.S (2019/09490-7). C.R.S.G. is supported by “bolsa de produtividade” by CNPq. This work was supported by the National Institutes of Health grant number R01-AI125565 to M.L. and T32-GM125592 to V.A.B. We are grateful to Haemocentro Hospital do Servidor Público for providing blood and plasma. Work in C.D. laboratory is supported by RMIT University and by grant APP2003712 from the Australian Health and Medical Research Council (NHMRC). We thank Prof. Paolo Di Mascio and Graziella E. Ronsein for the Mass spectrometry analyses performed at the Redox Proteomics Core of the Mass Spectrometry Resource at Chemistry Institute, University of São Paulo (FAPESP 2012/12663-1, 2016/00696-3, 2023/00995-4, CEPID Redoxoma 2013/07937-8), and Dr. Mariana P. Massafera for her technical assistance.

## Additional Information

Additional figures and tables are provided separately in Supplemental Figures and Supplemental Tables.

## Supplemental Figures

**Supplemental Figure 1.** **(A)** Coomassie-stained 6% SDS‒PAGE gel showing the parasite lysate of wild-type 3D7 and *Pf*MORC^GFP^ after coimmunoprecipitation with anti-GFP magnetic beads. Both lanes were used for mass spectrometry analysis. **(B)** Histogram shows the total proteins identified in mass spectrometry analysis from three biological replicates in wild-type 3D7 and *Pf*MORC^GFP^ coimmunoprecipitated samples. Venn diagram illustrates the labeled free LC-MS/MS enrichment of peptide hits obtained from **(C)** 3D7 control and **(D)** from *Pf*MORC^GFP^ parasites lysate. Briefly, 32 hpi (±4 h) trophozoite stage parasites were harvested and lysed, followed by incubation with anti-GFP-Trap-A beads from three independent biological replicates were used for quantification. False discovery rate (FDR) of 1% and peptides ≥2 leads to identifying 191, 814, 589 and 211, 617, 656 significant proteins in 3D7 and *Pf*MORC^GFP^, respectively. **(E)** MS/MS normalization of identified proteins from 3D7 parasites expressing *Pf*MORC and transgenic parasites expressing GFP (*Pf*MORC^GFP^) was carried out. Gene ontology classification showing biological process, cellular component, and molecular function of *Pf*MORC^GFP^/3D7 normalized proteins showing fold change ≥ 1.5.

**Supplemental Figure 2.** **(A)** Correlation plot (DeepTools PlotCorrelation) of the 30 h samples compared to the negative control ChIP-seq sample. **(B)** Correlation plot (DeepTools PlotCorrelation) of the 40 h samples compared to the negative control ChIP-seq sample. **(C)** Violin plot showing the ChIP-seq fold enrichment values of significantly called peaks in all 6 biological replicates. The two GFP samples were only used as additional controls for comparison purposes. **(D)** Venn diagram comparing the overlap of MACS2-called peaks between anti-HA biological replicates at 30 h. **(E)** Venn diagram comparing the overlap of MACS2-called peaks between anti-HA biological replicates at 40 h. **(F)** Venn diagram comparing the overlap of MACS2-called peaks between anti-GFP biological replicates at 40 h.

**Supplemental Figure 3.** **(Top)** Profile plot of the mean *Pf*MORC ChIP-seq fold enrichment (Log2[IP/Input]) for all four samples across all PfEMP1 (var) gene 5’ upstream regions and gene bodies. **(Bottom)** Heatmap of the *Pf*MORC ChIP-seq fold enrichment (Log2[IP/Input]) for all four samples across all *Pf*EMP1 (*var*) gene 5’ upstream regions and gene bodies. **(Inset to the right)** Zoom in on the average enrichment of *Pf*MORC at var genes with annotated exons.

**Supplemental Figure 4.** **(Top)** Profile plot of the mean *Pf*MORC ChIP-seq fold enrichment (Log2[IP/Input]) for all four samples across all *rif* gene 5’ upstream regions and gene bodies. **(Bottom)** Heatmap of the *Pf*MORC ChIP-seq fold enrichment (Log2[IP/Input]) for all four samples across all *rif* gene 5’ upstream regions and gene bodies. **(Inset to the right)** Zoom in on the average enrichment of *Pf*MORC at *rif* genes with annotated exons.

**Supplemental Figure 5.** **(Top)** Profile plot of the mean *Pf*MORC ChIP-seq fold enrichment (Log2[IP/Input]) for all four samples across all *rif* gene 5’ upstream regions and gene bodies. **(Bottom)** Heatmap of the *Pf*MORC ChIP-seq fold enrichment (Log2[IP/Input]) for all four samples across all *rif* gene 5’ upstream regions and gene bodies. **(Inset to the right)** Zoom in on the average enrichment of *Pf*MORC at *rif* genes with annotated exons.

**Supplemental Figure 6.** The heatmaps show the transcript abundance (Chappell et al. 2020) of putative *Pf*MORC gene targets at 30 h (left) and 40 h (right). Red signifies high transcript abundance, and green signifies low transcript abundance. Both timepoints are organized into two major clusters (highlighted with the yellow bar and blue bar).

**Supplemental Figure 7.** **(A)** Associated with Figure 3A,B. Mean fold enrichment (Log2[IP/Input]) summary plot (top) and full heatmap (bottom) of fold enrichment of PfMORC, six associated ApiAP2 factors (AP2-G5, AP2-O5, AP2-I, PF3D7_1107800, PF3D7_0613800, and PF3D7_1239200), HP1, and a negative no-epitope control across *Pf*MORC binding sites at the 30 h and 40 h timepoints. **(B)** Quantitative Venn diagrams of the binding site overlaps between *Pf*MORC and the six associated ApiAP2 factors.

**Supplemental Figure 8.** DNA motif analyses from these different categories: (1) Unique to 30 hpi ChIP-seq Timepoint, (2) 30hpi ChIP-seq Timepoint, (3) Overlap of ChIP-seq Timepoints, (4) 40hpi ChIP-seq Timepoint, and (5) Unique to 30hpi ChIP-seq Timepoint. The values to the right of each motif contain the enrichment value, number of peaks containing that motif, and percent of the peaks the contain that motif calculated by Meme Suite.

**Supplemental Figure 9.** Mean fold enrichment (Log2[IP/Input]) summary plot **(top)** and full heatmap **(bottom)** of fold enrichment of ten selected epigenetic marks (H2A.Z, H3K9ac, H3K4me3, H3K27ac, H3K18ac, H3K9me3, H3K36me2/3, H4K20me3, and H3K4me1) across *Pf*MORC binding sites at the 30 h and 40 h timepoints.

**Supplemental Figure 10.** Comparison of transcriptional changes with melatonin treatment. **(A)** Volcano plot showing the differentially expressed genes in *Pf*MORC-KD parasites relative to *Pf*MORC-WT after 100 nM melatonin treatment for 5h from three independent experiments. **(B)** Venn diagram shows intersecting DEGs from the experiment with KD vs WT with DEGs obtained from the experiment (KD vs WT) treated with 100 nM melatonin for 5 h. Number of DEGs are shown as up (red) and downregulated (green). The intersecting region shows 282 upregulated and 340 downregulated genes. Heatmap showing significant DEGs based on *p-values* and log_2_FD for upregulating **(C)** and downregulating **(D)**. These genes are taken from 622 intersecting DEGs showing partial changes in expression after melatonin treatment.

