## Supplementary figures and images for "A *Plasmodium falciparum* MORC protein complex modulates epigenetic control of gene expression through interaction with heterochromatin"

### Supplemental Figure 1

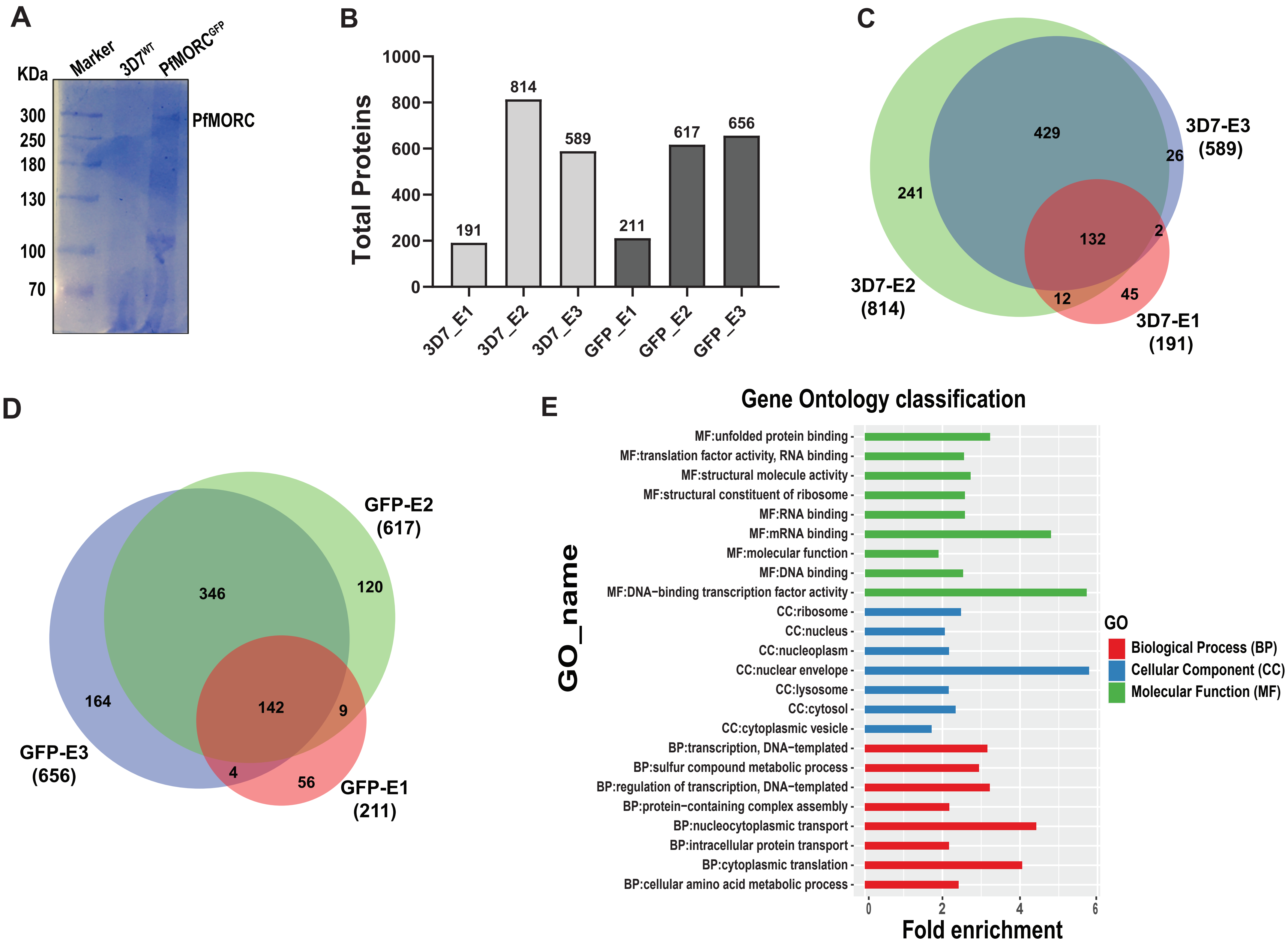

### Supplemental Figure 2

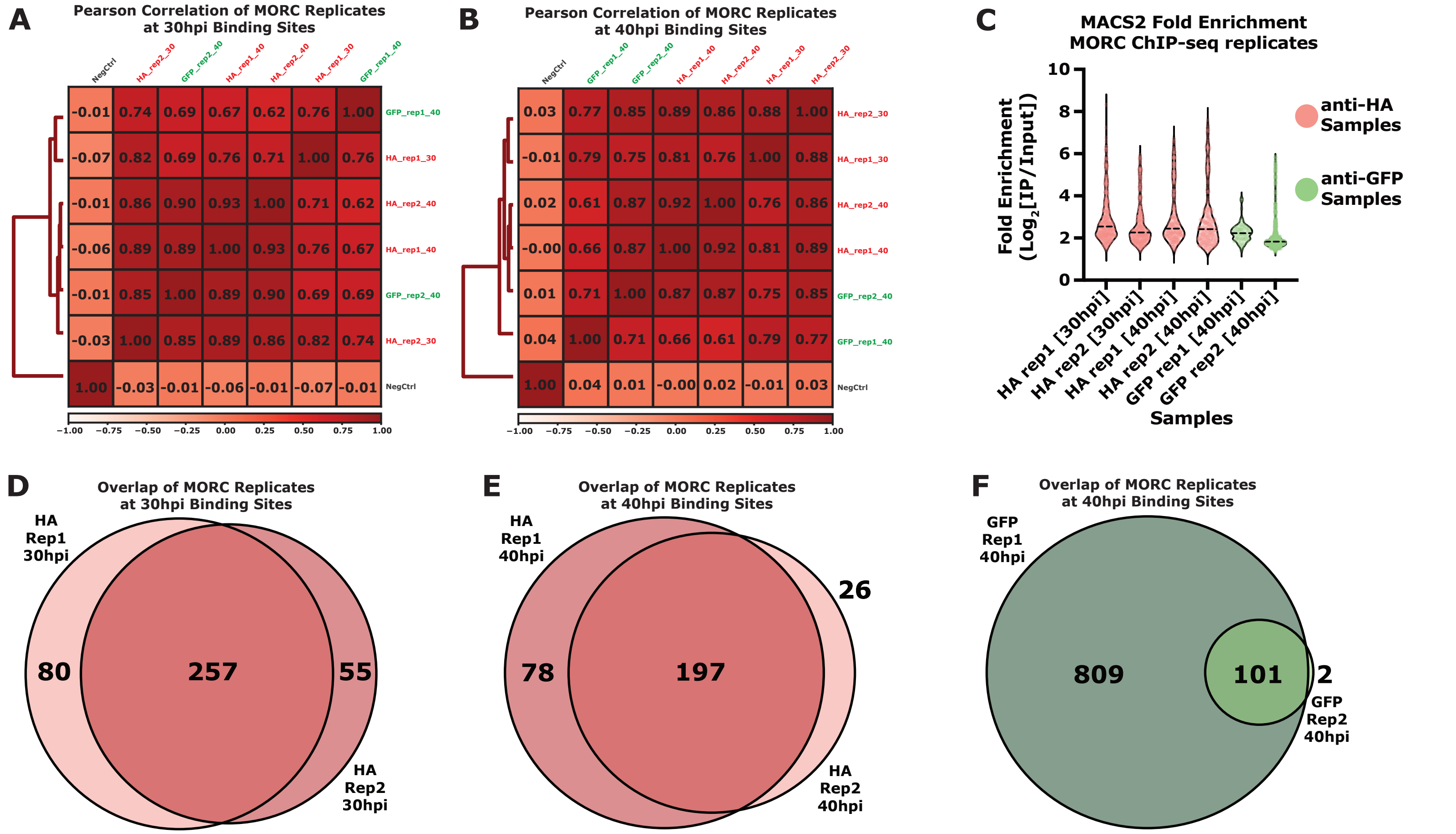

### Supplemental Figure 3

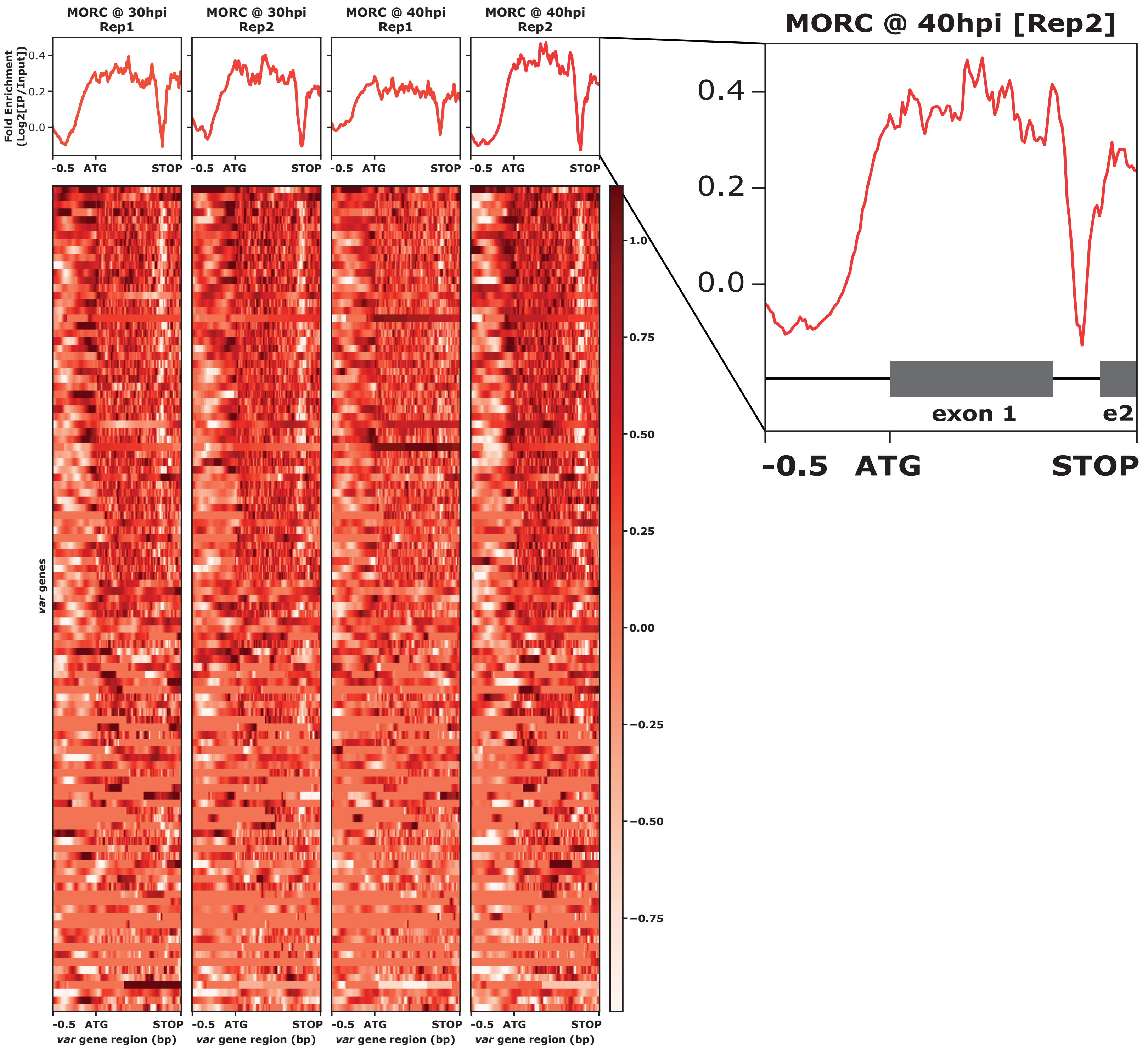

### Supplemental Figure 4

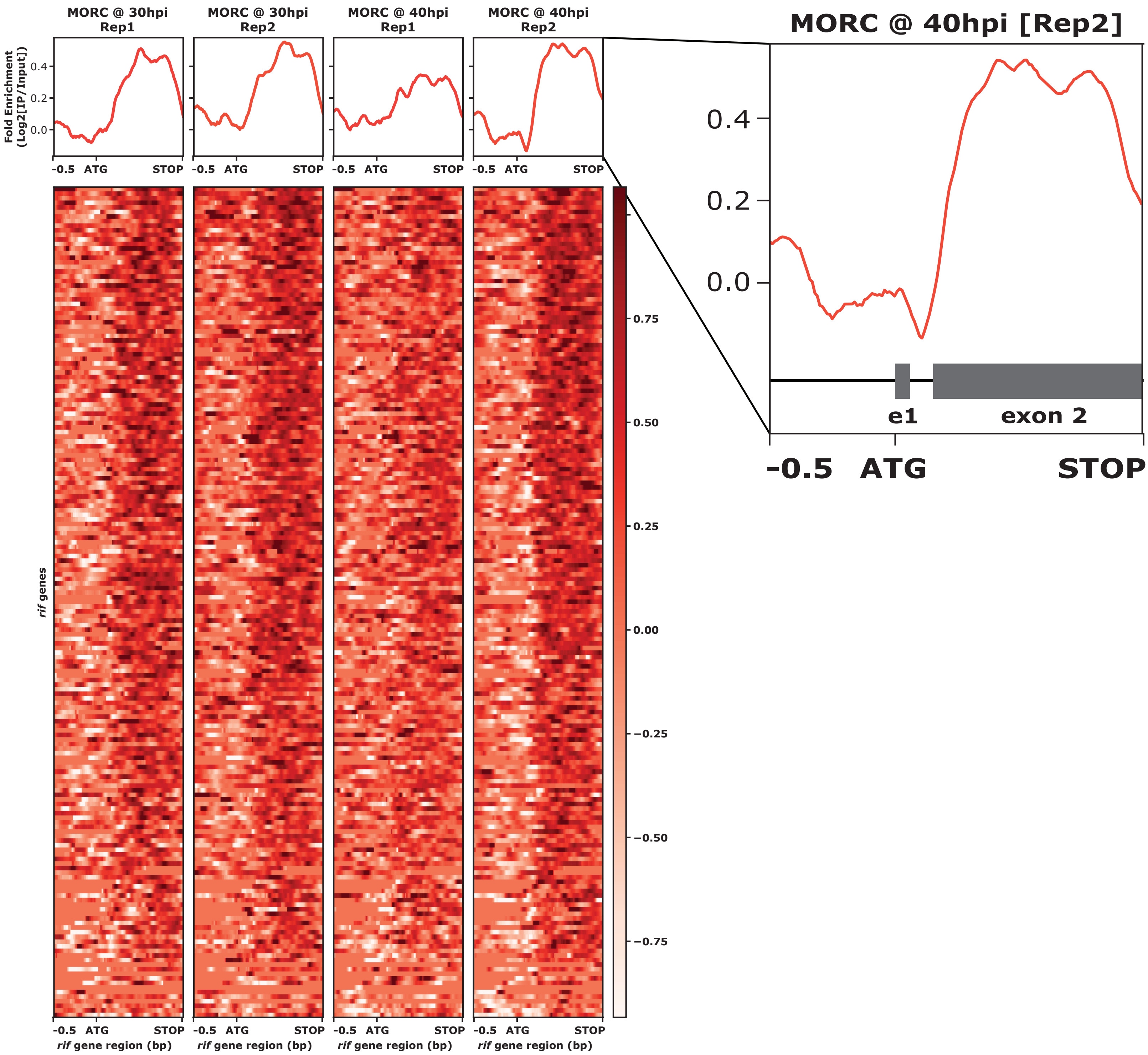

### Supplemental Figure 5

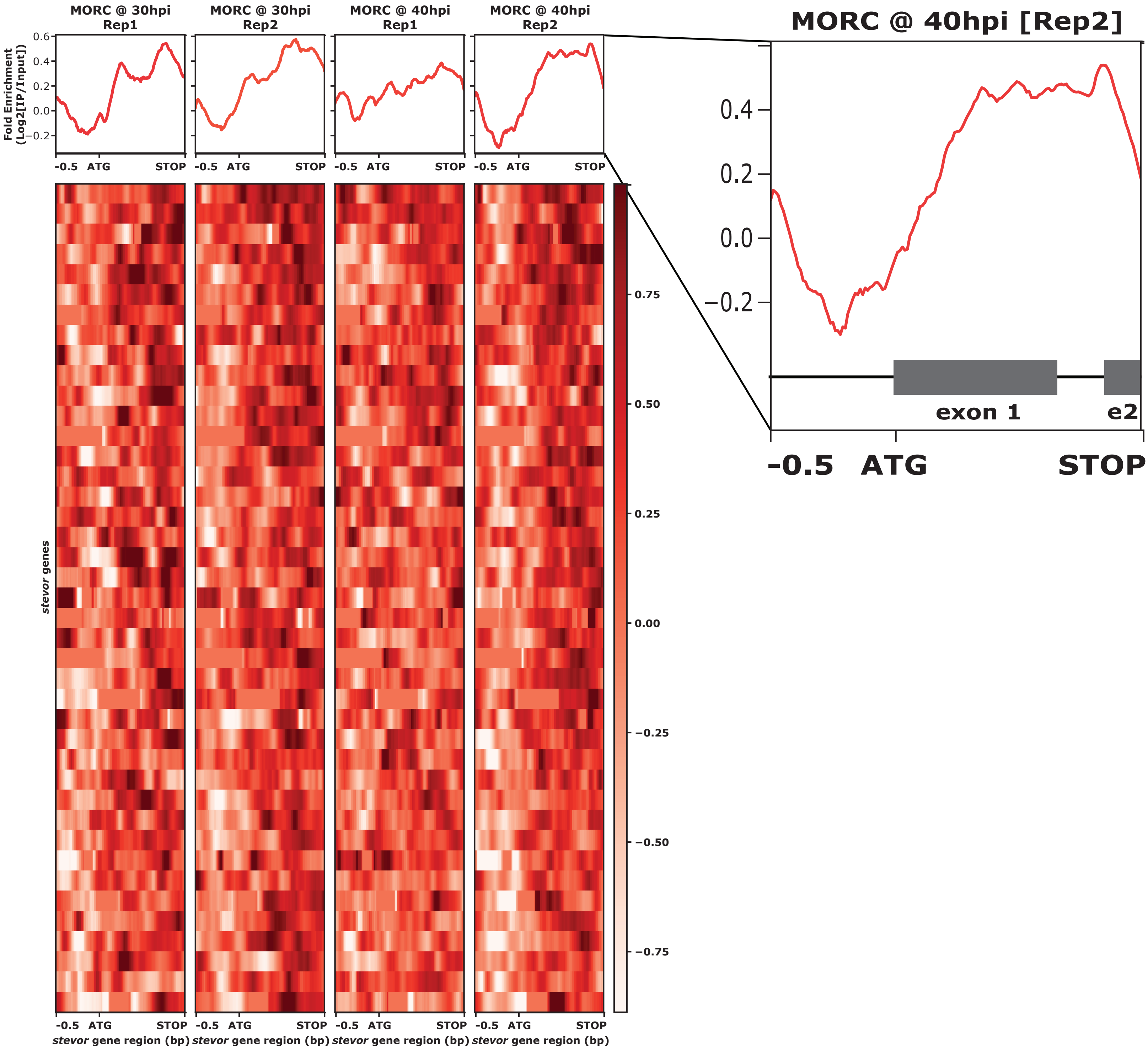

### Supplemental Figure 6

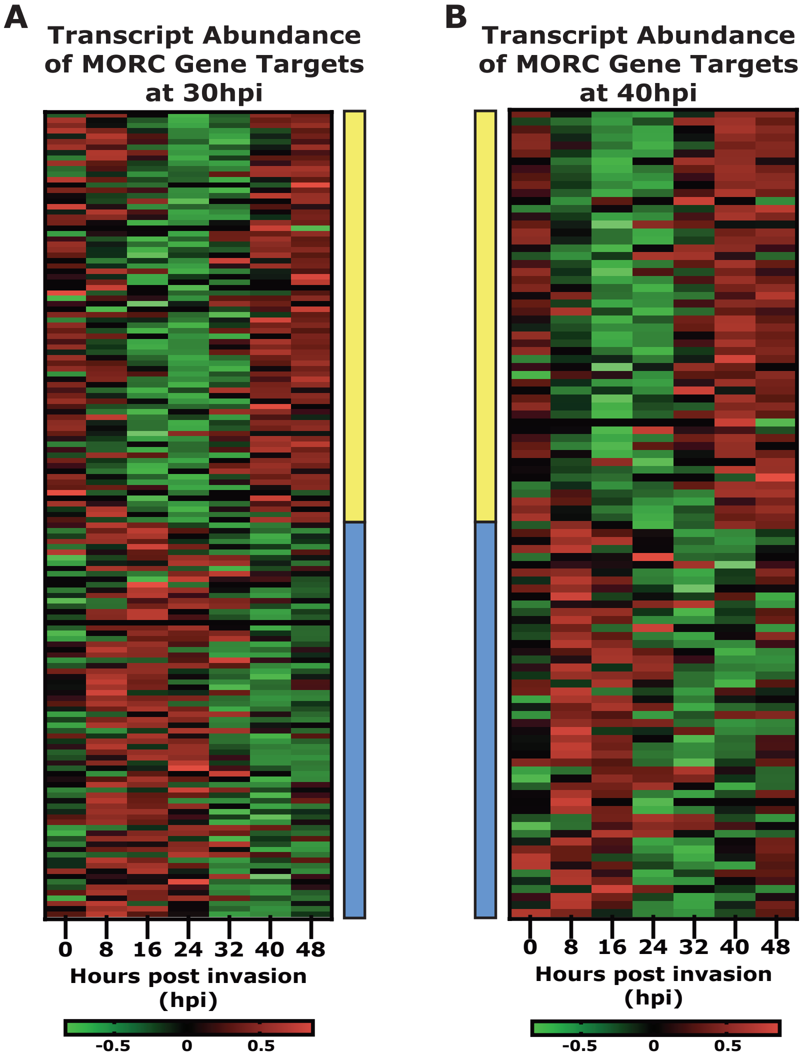

### Supplemental Figure 7

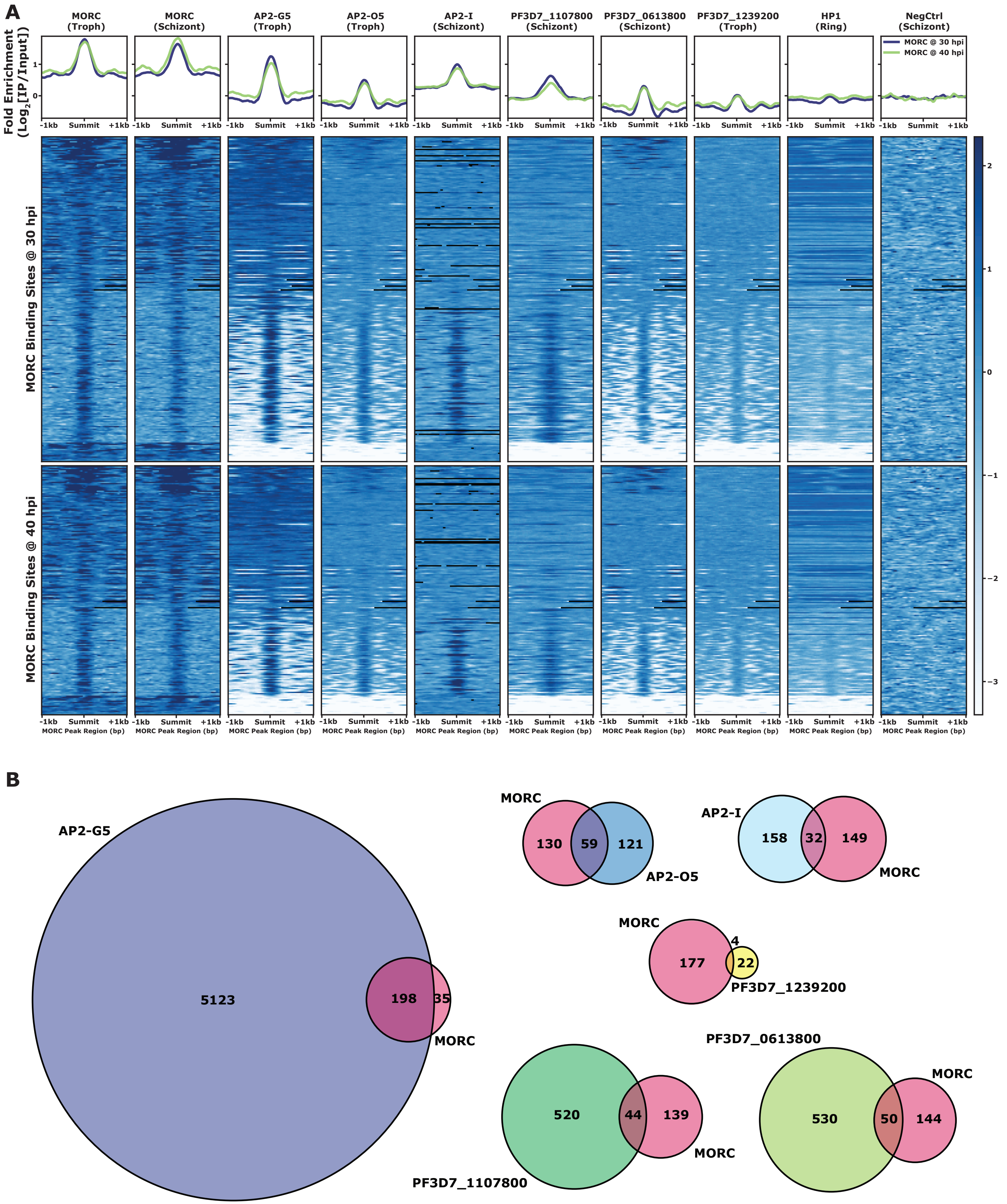

### Supplemental Figure 8

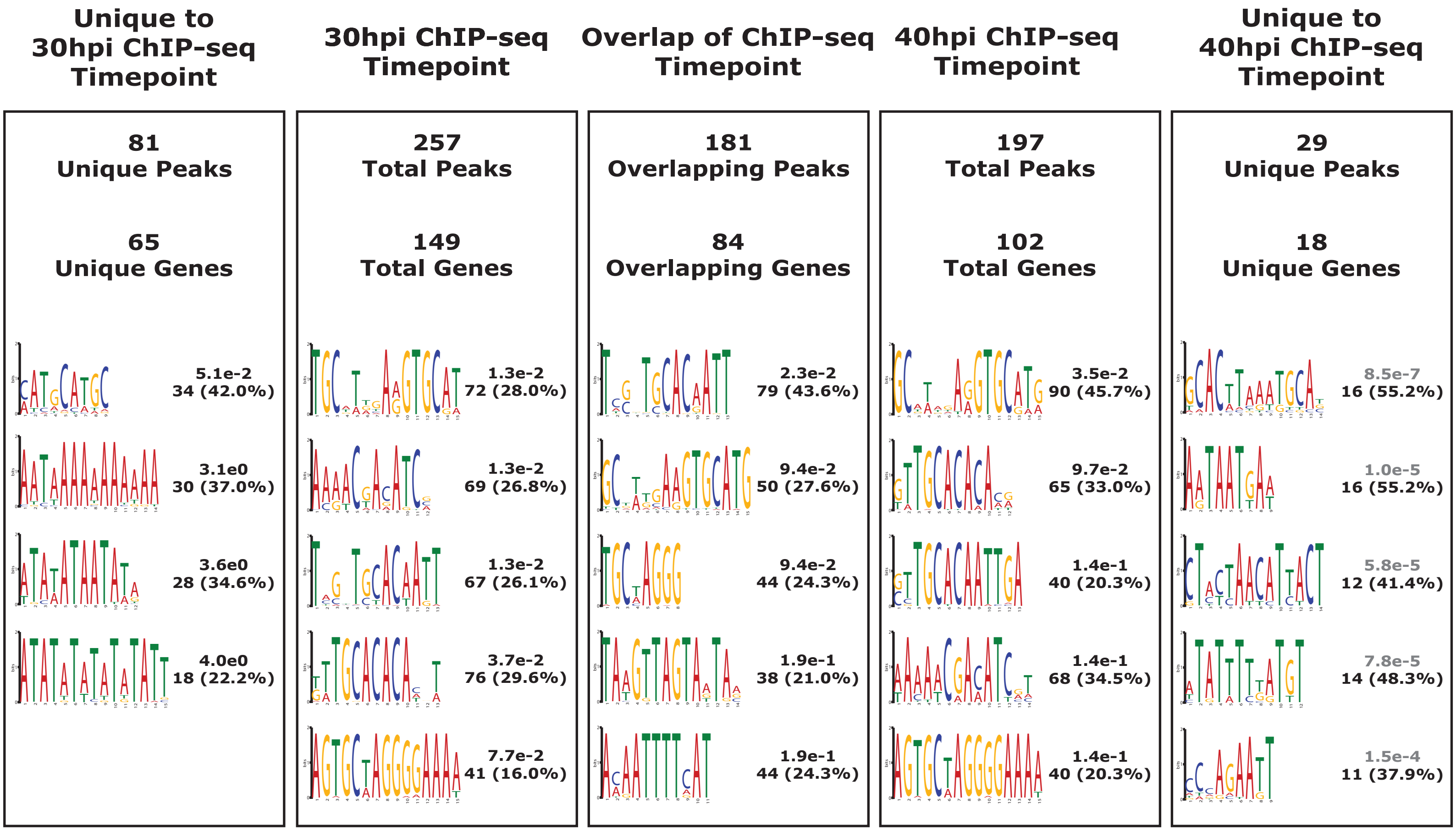

### Supplemental Figure 9

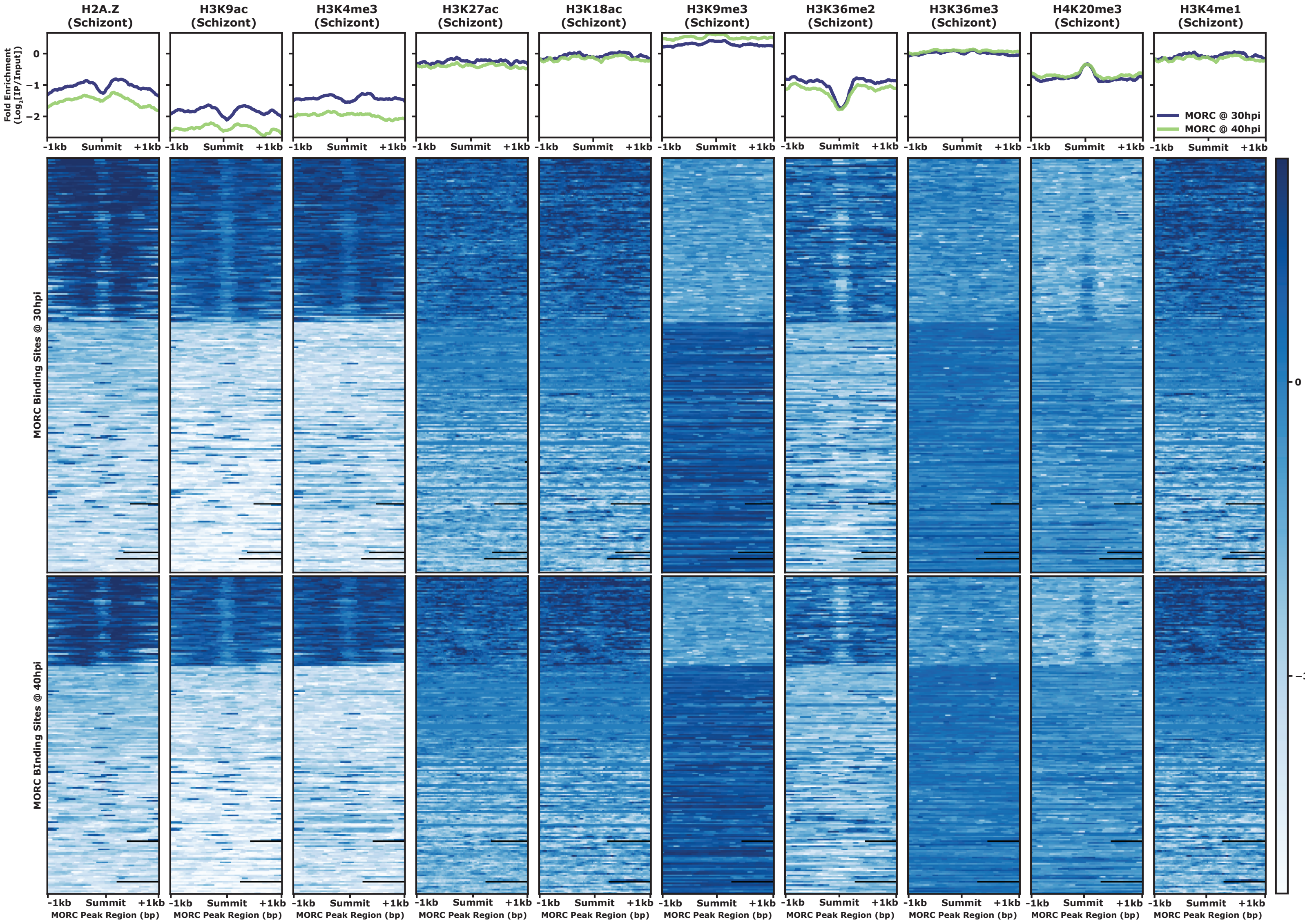

### Supplemental Figure 10

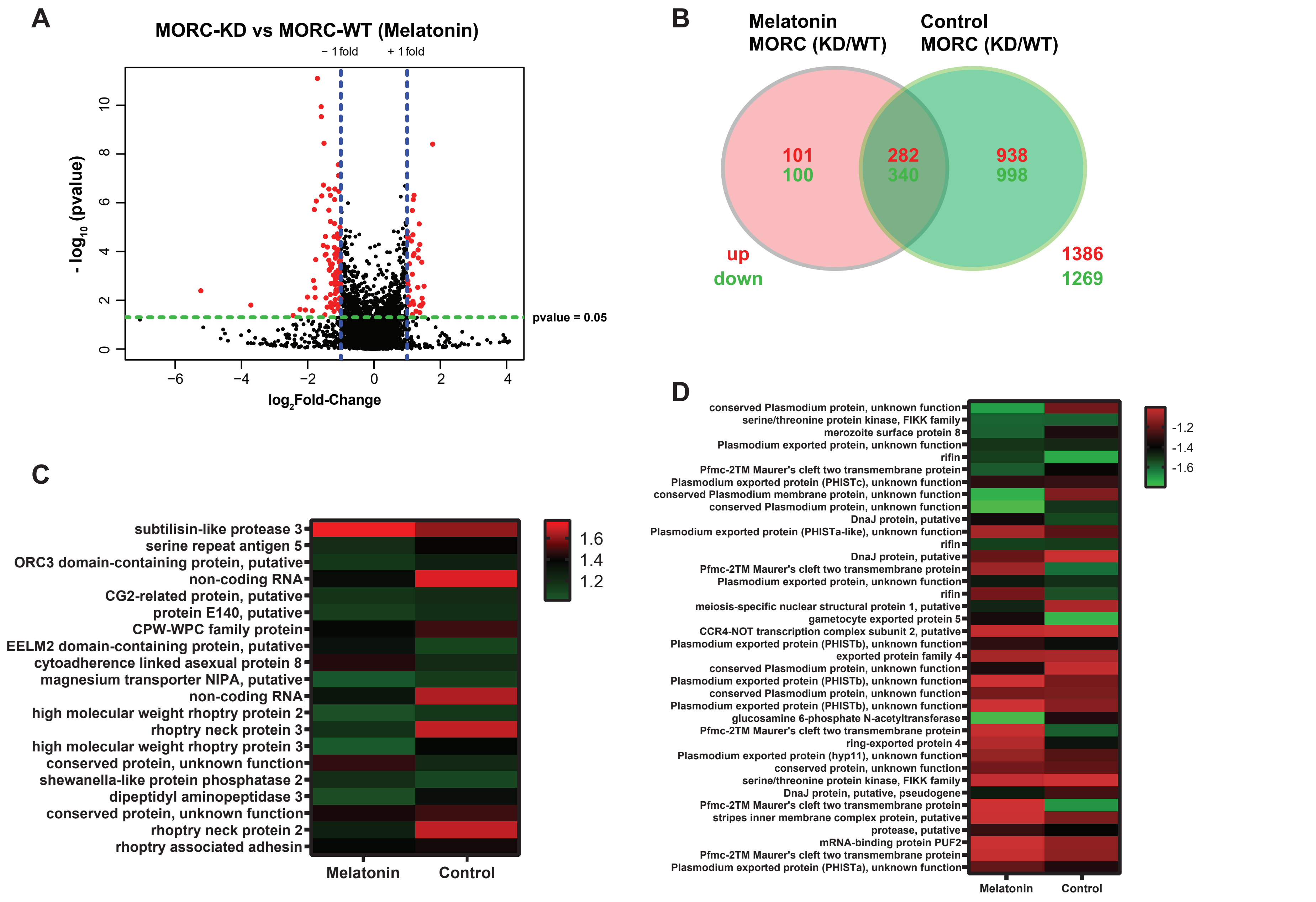
